# The RNA-binding domain of hnRNP U extends beyond the RGG/RG motifs

**DOI:** 10.1101/2023.09.20.558674

**Authors:** Otto A. Kletzien, Deborah S. Wuttke, Robert T. Batey

**Affiliations:** Department of Biochemistry, University of Colorado, Boulder, CO 80309-0596, USA

**Author notes:** To whom correspondence should be addressed. DSW. RTB.

## Abstract

Heterogeneous nuclear ribonucleoprotein U (hnRNP U) is a ubiquitously expressed protein that regulates chromatin architecture through its interactions with numerous DNA, protein, and RNA partners. The RNA-binding domain (RBD) of hnRNP U was previously mapped to an RGG/RG element within its disordered C-terminal region, but little is understood about its binding mode and potential for selective RNA recognition. Analysis of publicly available hnRNP U enhanced UV crosslinking and immunoprecipitation (eCLIP) data identified high-confidence binding sites within human RNAs. We synthesized a set of diverse RNAs encompassing eleven of these identified crosslink sites for biochemical characterization using a combination of fluorescence anisotropy and electrophoretic mobility shift assays. These *in vitro* binding experiments with a rationally designed set of RNAs and hnRNP U domains revealed that the RGG/RG element is a small part of a more expansive RBD that encompasses most of the disordered C-terminal region. This RBD contains a second, previously experimentally uncharacterized RGG/RG element with RNA-binding properties comparable to the canonical RGG/RG element. These RGG/RG elements serve redundant functions, with neither serving as the primary RBD. While in isolation each RGG/RG element has modest affinity for RNA, together they significantly enhance the association of hnRNP U with RNA, enabling binding of most of the designed RNA set with low to mid-nanomolar binding affinities. Identification and characterization of the complete hnRNP U RBD highlights the perils of a reductionist approach to defining biochemical activities in this system and paves the way for a detailed investigation of its RNA-binding specificity.

## INTRODUCTION

Heterogeneous nuclear ribonucleoprotein U (hnRNP U, also known as scaffold attachment factor A or SAF-A) is a ubiquitously expressed, predominantly nuclear protein that regulates chromatin architecture through its interactions with numerous RNA, DNA and protein binding partners.^1–3^ Because it contains a DNA-binding SAP domain, an SPRY/B30.2 protein-protein interaction domain, a AAA+ ATP-dependent oligomerization domain and a C-terminal domain containing two RNA-binding RGG/RG elements, hnRNP U can act as a bridge between various components of the nuclear machinery (**Figure 1**).^3–4^ Previous studies have implicated hnRNP U as a key component of the nuclear scaffold, a dynamic RNA-protein network that maintains chromatin architecture.^1–3^ Within the nuclear scaffold, oligomerization of hnRNP U through its AAA+ domain is required to maintain open chromatin, and this oligomerization only occurs when the neighboring RGG/RG element is bound to RNA.^3, 5^ Apart from its scaffolding functions, hnRNP U mediates numerous other cellular processes through its interactions with RNA, such as tethering the *Xist* lncRNA to the inactive X chromosome,^4^ removing RNA from DNA damage sites to prevent aberrant DNA repair,^6^ promoting antibody diversity by binding R-loops and promoting NHEJ-mediated DNA repair at the IgH gene locus,^7^ and regulating maturation of U2 snRNA by inhibiting trafficking of the 12*S* U2 precursor RNA into Cajal bodies.^8^

**Figure 1:**
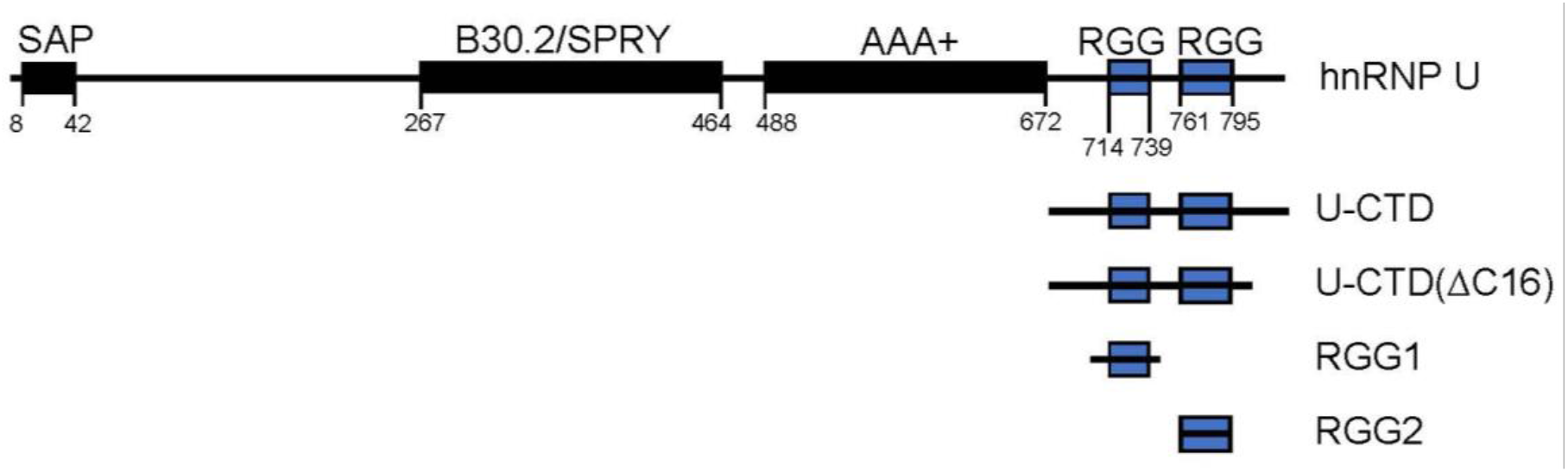
Domain map of human hnRNP U. The full length protein (top) contains a C-terminal RNA binding domain (U-CTD) containing two RGG/RG motifs (blue boxes). Included in this figure are the fragments of U-CTD that were used in this study.

The RNA-binding activity of hnRNP U^8–15^ is proposed to be mediated by a single RGG/RG element spanning amino acids 715-739 (RGG1, **Figure 1**).^16^ However, the bioinformatic identification of a potential second RGG/RG element (RGG2, **Figure 1**) within the disordered CTD of hnRNP U suggests that the canonically defined RGG/RG element may not represent the full RNA-binding domain (RBD) of hnRNP U.^17^ The RGG/RG element (or RGG/RG motif), a type of low-complexity region defined by arginine- and glycine-rich repeats, is one of the most common RBDs in the human proteome, found in over 1,000 human proteins.^17–18^ RGG/RG elements can be categorized as Di-RGG, Tri-RGG, Di-RG, or Tri-RG, depending on the number of repeats separated by ≤4 amino acids and the number of glycines within each repeat,^17^ although it remains unknown whether RG- and RGG-regions possess differing RNA-binding properties. Despite their abundance and functional importance, development of a structural basis for RNA recognition by RGG/RG elements has been difficult due to their intrinsically disordered nature in the free state and likely multiple and heterogeneous RNA recognition modes in the bound state.^19^ Because of this lack of detailed structural information, the protein sequence features driving affinity and selectivity for RNA require biochemical definition. In addition to arginine and glycine, RGG/RG elements are commonly enriched in aromatic amino acids, suggesting a potential role for π-stacking interactions in RNA recognition.^19–20^ Since the few available structures of RGG/RG–RNA complexes have not included aromatic amino acids,^21–22^ the specific contribution of this element to RNA recognition unknown.

Unlike the majority of RGG/RG containing proteins, hnRNP U contains no other identified RBDs adjacent to the RGG/RG element,^17–18^ making it an excellent model system to study RGG/RG-mediated RNA recognition. However, despite the biological importance of its many interactions with RNA, biochemical data on the RNA-binding preferences of hnRNP U remains incomplete and contradictory. Previous crosslinking and immunoprecipitation (CLIP) studies have revealed that hnRNP U binds to a large number of RNAs, including virtually every class of small ncRNA,^8, 23^ but these interactions have not been biochemically defined. Earlier studies of the RNA-binding capabilities of the canonical RGG/RG element of hnRNP U were performed using polyribonucleotide homopolymers, which are unlikely to be representative of biologically relevant RNAs.^16^ A large C-terminal hnRNP U fragment containing both RGG/RG motifs was shown to bind an array of RNA ligands with low affinity for simple hairpins, ssRNA, and poly-A RNA, and with much higher affinity for RNAs containing complex secondary structures.^9^ However, this study focused almost exclusively upon non-biological RNA substrates of RGG/RG elements and therefore the relevance of these conclusions regarding binding selectivity to biological RNAs remains unknown. Recently, a study of the role of hnRNP U and hnRNP L in alternative splicing of *MALT1* pre-mRNA found that hnRNP U associates with GU-containing hairpins within the message and stabilizes secondary structure.^24^

Here, we characterize the RNA-binding domain of human hnRNP U in depth using a panel of biologically relevant RNA ligands to enable a detailed investigation of its RNA-binding properties in a biologically relevant context. High-confidence RNA binding sites were identified from publicly available eCLIP data and RNA fragments modeled upon eleven of these sites were synthesized for *in vitro* binding assays with purified hnRNP U. Full-length hnRNP U binds these RNA fragments with a wide range of affinities spanning 11 – 2200 nM. A protein construct containing the full, disordered C-terminal domain (U-CTD) recapitulates the binding affinity of the full-length protein, confirming that hnRNP U binds RNA through this domain. Using a series of protein fragments spanning the C-terminal region of hnRNP U, the boundaries of the RBD were more precisely defined. We found that the full RBD covers the entire disordered C-terminal domain, including the bioinformatically identified RGG2 element, which fits the description of a “Tri-RG”.^17^ In isolation, the RGG/RG elements exhibit similar RNA-binding preferences, but RGG2 binds with approximately two-fold weaker affinity than RGG1, suggesting that the different number of RG and RGG repeats may tune their ability to bind RNA. Further, we show that the full RBD can tolerate the loss of either RGG/RG motif with only slight losses of affinity, indicating that neither is essential for full RNA binding activity. Rather, high-affinity binding is driven by largely redundant RGG/RG elements working in concert with their flanking sequences. The newly observed binding contributions of the regions flanking both RGG/RG elements suggest that prior studies using minimized RGG/RG element constructs might miss important contextual features that affect the RNA binding properties of native proteins.

## Materials and Methods

### Bioinformatic analysis of hnRNP U binding sites

Human hnRNP U eCLIP data in K562 and HepG2 cell lines was downloaded from the ENCODE Consortium database^25^ (experiment ENCSR240MVJ: files ENCFF587PLY, ENCFF125EPG, ENCFF027WDY; experiment ENCSR520BZQ: files ENCFF025YVA, ENCFF197OBL, ENCFF188FGR). Each of the two biological replicates was used as input for PureCLIP^26^ with the following settings: chromosomes 1-5 used to train model, ENCODE input alignments ENCFF027WDY and ENCFF188FGR used as a control, merge distance = 8, and bc_1 protein characteristic. For each cell line, bedtools intersect^27^ was used to identify PureCLIP binding regions conserved across both biological replicates. The final hnRNP U binding regions of each cell line were then merged with bedtools intersect, yielding the final crosslink sites conserved across both cell lines. To ensure proper normalization over background, bedtools multicov^27^ was used to quantify eCLIP reads over each hnRNP U crosslink site. Briefly, we averaged the number of reads in both biological replicates of each cell line, then subtracted the number of reads in the corresponding input. We then averaged both cell lines together and regions with an average enrichment of ≤0 were removed from the final pool. The resulting 323 crosslink sites were annotated according to human genome assembly GRCh38. Crosslink sites were ranked according to the mean PureCLIP score across all four biological replicates.

To identify sequence motifs, the final hnRNP U-bound regions were extended by four nucleotides (nt) in each direction using bedtools slop and used as input for MEME^28^ with the following settings: any number of repeats (anr), 1st-order background and motif minimum width = 4 nt. Because the larger HepG2 and K562 PureCLIP datasets contained too many sites for accurate motif detection by MEME, motifs in these datasets were detected using STREME with the following settings: 1st-order background, 0.05 p-value threshold, motif minimum width = 4 nt.^29^ To verify that the selected survey RNAs represent bona fide peaks, the raw eCLIP alignments from ENCODE were visualized in the Integrated Genomics Viewer (IGV) and compared with the corresponding input alignments.^30^

### RNA secondary structure prediction

For all synthesized RNAs, secondary structures were initially predicted using Mfold software (http://www.unafold.org/mfold/applications/rna-folding-form.php).^31^ The following default conditions were used for all structure predictions: RNA is specified to be linear, folding temperature = 37 °C, ionic conditions = 1 M NaCl, suboptimality percent = 5, upper bound on number of computed foldings = 50, window parameter = default, maximum interior/bulge loop size = 30 nt, maximum asymmetry of an interior/bulge loop = 30, and no limit on the maximum distance between paired bases.

### Expression and purification of proteins

All human hnRNP U protein (Isoform 2, which lacks the disordered region spanning amino acids 218-231; Uniprot Q00839) variants were cloned into the pHMM expression vector^32^ and expressed as N-terminal 8xHis-MBP fusions. Sequence-verified expression vectors were transformed into *E. coli* Rosetta (DE3) pLysS cells, plated on 2xYT plates supplemented with 34 μg/mL chloramphenicol and 50 μg/mL kanamycin and incubated at 37 °C overnight. An individual colony was transferred to 4 mL 2xYT liquid culture grown overnight at 37 °C in the presence of chloramphenicol and kanamycin. For large-scale cultures, 1 mL of the initial overnight culture was used to inoculate 1 L of 2xYT medium supplemented with 34 μg/mL chloramphenicol and 50 μg/mL kanamycin. Cultures were grown at 37 °C until reaching an OD_600_ of 0.4-0.6, at which time each culture was placed on ice for 5 minutes, induced with 0.5 mM IPTG and continued to incubate at 25 °C for 16 to 20 hours.

To harvest cells, cultures were centrifuged at 5,000 x*g* for 30 minutes and the cell pellets resuspended in Lysis Buffer (1 M KCl, 50 mM Tris pH 7.4, 10 mM imidazole, 2 mM MgCl_2_, 10% glycerol). To disfavor co-purification of nucleic acids from *E. coli*, all buffers used in the Ni-NTA purification contained 1 M KCl. Cell lysis was performed using an Avestin Emulsiflex C3 homogenizer (ATA Scientific). Cell lysates were clarified by additional centrifugation at 14,000 x*g* for 30 minutes and supernatants were incubated 1 hour at 4 °C with 5 mL of His-Pur™ Ni-NTA resin (Qiagen) previously washed 3 times with 25 mL of Lysis Buffer. After incubation, resin was centrifuged at 1,500 x*g* for three minutes, supernatant was removed, and resin was washed with five volumes of Lysis Buffer. The centrifugation and wash steps were repeated one time with five volumes of Wash Buffer (1 M KCl, 50 mM Tris pH 7.4, 100 mM imidazole, 2 mM MgCl_2_, 10% glycerol). After the second wash, the resin was incubated with five volumes of Elution Buffer (1 M KCl, 50 mM Tris pH 7.4, 350 mM imidazole, 2 mM MgCl_2_, 10% glycerol) for 10 minutes at 4 °C, then centrifuged as before. The eluate was collected and concentrated down to ∼3 mL in 10 kDa molecular weight cutoff centrifugal concentrator tubes (Amicon). The concentrated eluate was loaded onto a size exclusion column (SEC) (HiLoad 16/600 pg75 or pg200; GE Life Sciences) equilibrated with SEC buffer (300 mM KCl, 100 mM Tris pH 7.4, 2 mM MgCl_2_, 2 mM DTT, 10% glycerol) and run at a flow rate of 1 mL/min for 1.3 column volumes. SEC fractions were analyzed by SDS-PAGE (**Figure S1**), pooled, concentrated to >100 µM, aliquoted, frozen in liquid nitrogen, and stored at −20 °C.

### Preparation, purification, and 3’ FTSC labeling of RNAs

dsDNA templates for use in *in vitro* transcription reactions with T7 RNA polymerase were assembled from ssDNA primers (Integrated DNA Technologies) using 30 cycles of recursive PCR using a standard protocol with modified primer concentrations (1 μM 5’ and 3’ outer primers, 10 nM inner primers).^33^ 200-300 µL of each PCR reaction was used as template for 2-3 mL *in vitro* transcription reactions with T7 RNA polymerase and inorganic pyrophosphatase, followed by purification on denaturing PAGE gels as previously described.^34^ RNAs to be labeled with fluorescein were ethanol precipitated with 2.5 volumes of cold 100% ethanol, 0.1 volume of 3 M sodium acetate, pH 5.2, and 1 μL of glycogen followed by storage at −20 °C, whereas all other RNAs were simply frozen at −20 °C.

For use in fluorescence-based experiments, RNAs were labeled at the 3’-end with fluorescein thiosemicarbazide (FTSC), as previously described.^35^ Following the labeling reaction, RNAs were pelleted by centrifugation at 16,100 x*g* for 15 minutes at 4 °C, then pellets were resuspended in 100 μL Milli-Q H_2_O, denatured with >200 μL formamide, incubated at 65 °C for >5 minutes and purified by electrophoresis on a denaturing acrylamide gel ranging from 6-15% depending on the size of the RNA, run in 1x TBE electrophoresis running buffer. RNA bands were readily visible due to the fluorescein label and were excised with a razor blade, then extracted through overnight crush-and-soak in ∼900 μL Milli-Q H_2_O. The eluate was passed through a 0.22 μm Corning Co-Star microfuge filter tube and the filtrate was transferred to an Amicon Ultra microfuge concentrator tube (3 or 10 kDa MWCO), concentrated to ∼50 μL, washed twice with ∼300 μL Milli-Q H_2_O, and concentrated to a final volume <50 µL. RNA concentrations were calculated using the molar extinction coefficients calculated by IDT OligoAnalyzer and measured absorbance at 260 nm.

### Fluorescence Anisotropy (FA) Measurements

Prior to each experiment, the FTSC-labeled RNAs were annealed by a snap-cool method in which the RNA was heated to 85 °C for 3 minutes and then immediately placed on slushy ice for at least 3 minutes. Snap-cooled RNAs were diluted to 8 nM in 2x RNA master mix (0.2 mg/mL bovine serum albumin (molecular biology grade, NEB), 0.2 mg/mL yeast tRNA, 10% glycerol) at room temperature. Protein aliquots were thawed, diluted in SEC buffer, centrifuged at 16,100 x*g* for 15 minutes at 4 °C, and supernatant was transferred to a new tube. Protein concentrations were measured by 280 nm absorbance, then diluted to double the starting concentration of the titration. Serial dilutions were prepared in SEC buffer at room temperature. In a flat-bottom low-flange 384-well black polystyrene plate (Corning, #3575), 10 µL of each protein serial dilution was added to 10 µL of 2x RNA master mix for a final volume of 20 µL containing 4 nM labeled RNA probe, gently mixed and incubated in the dark at room temperature for >45 minutes, to allow the reaction to reach equilibrium. The final buffer conditions were 150 mM KCl, 50 mM Tris pH 7.4, 1 mM MgCl_2_, 1 mM DTT, 10% glycerol, 0.1 mg/mL BSA, 0.1 mg/mL yeast tRNA, 4 nM FTSC-labeled RNA.

Using a ClarioStar Plus FP plate reader (BMG Labtech) fluorescence measurements were collected and converted to anisotropy using the equation

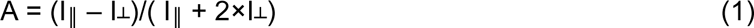

Where A is the fluorescence polarization, I_‖_ is the emission fluorescence intensity parallel to the excitation light and I_┴_ is the emission fluorescence intensity perpendicular to the excitation light. For each assay, 0.25 nM fluorescein was used to set the gain, to ensure consistent measurements between days. To determine the salt dependence of select interactions, the same conditions were set up as described but with final KCl concentrations ranging from 25 mM to 300 mM.

Equilibrium dissociation constants were determined by fitting in Kaleidagraph (Synergy Software) with either a simple one-transition Hill fit or a two-transition quadratic Hill fit, depending on the number of observed transitions in the binding isotherm. The one transition Hill fit equation used was

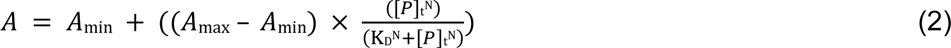

where A is anisotropy, A_min_ is the lower baseline, A_max_ is the upper baseline, [P]_t_ is the total protein concentration, N is the Hill coefficient (referred to as N_H_ elsewhere in text), and K_D_ is the apparent equilibrium dissociation constant. The two-transition quadratic Hill fit equation used was

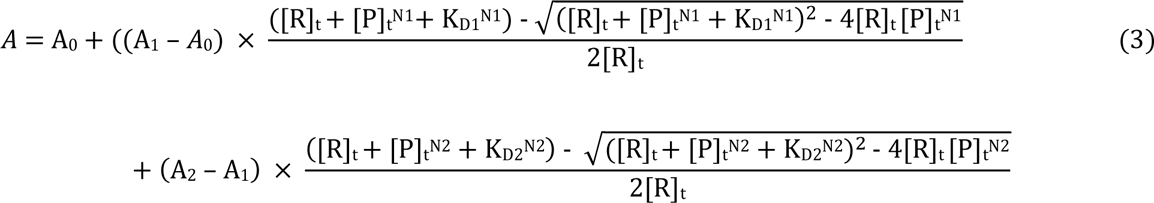

where A is anisotropy, A_0_ is the lower baseline, A_1_ is the upper baseline of the first transition, [R]_t_ is the total labeled RNA concentration, [P]_t_ is the total protein concentration, K_D1_ is the apparent equilibrium dissociation constant of the first transition, A_2_ is the upper baseline of the second transition, K_D2_ is the apparent equilibrium dissociation constant of the second transition, N1 is the Hill coefficient of the first transition, and N2 is the Hill coefficient of the second transition. To convert anisotropy to fraction bound, the relation

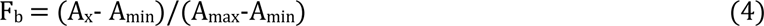

was used, where F_b_ is fraction bound, A_x_ is anisotropy, A_min_ is the lower baseline of the fitted curve, and A_max_ is the upper baseline of the fitted curve. For binding curves that did not appear to saturate at the highest protein concentrations, anisotropy values were normalized to other saturated curves that were run in parallel using the same protein serial dilution. Each reported K_D_ (apparent equilibrium dissociation constant) is the mean value of at least three separate technical repeats. The standard error of the mean (s.e.m.) was calculated for each K_D_ and reported.

### Electrophoretic Mobility Shift Assays (EMSAs)

RNA master mix and protein dilutions were prepared as above, with two differences: the reaction volume was halved (5 µL of RNA master mix and protein dilutions were added for a final volume of 10 µL), and the RNA was diluted to 24 nM in the 2x master mix for a final concentration of 12 nM. Reactions were incubated at room temperature for >45 minutes, then 8 µL was loaded onto a 6% 29:1 acrylamide:bisacrylamide gel in 0.5x TB running buffer (45 mM Tris-HCl, 45 mM borate, pH 8.2) while running at 5 W. Gels were run for 35-45 minutes, then imaged on a Typhoon imager (GE Healthcare) with 495 excitation and 520 emission wavelengths. Band intensities were quantified in ImageJ and used to calculate fraction bound. Dissociation constants were determined by fitting in Kaleidagraph with either a one- or two-transition quadratic Hill fit. The two-transition quadratic Hill fit was equation (2) described in the previous section, and the one-transition quadratic Hill fit is

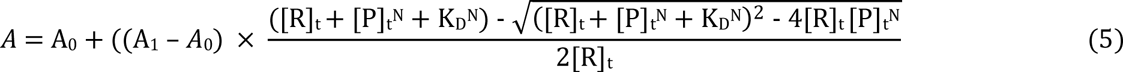

where A is anisotropy, A_0_ is the lower baseline, A_1_ is the upper baseline, [R]_t_ is the total labeled RNA concentration, [P]_t_ is the total protein concentration, K_D_ is the apparent equilibrium dissociation constant, and N is the Hill coefficient.

To measure the apparent dissociation constants of each individual bound band in EMSAs performed with Syncrip and MTRNR2L6 RNAs, the fraction bound was quantified in two ways. First, the total bound and unbound fractions were calculated by drawing a single box around all bound or unbound species to generate a macroscopic K_D_ value, as determined by a Hill fit in Kaleidagraph. Second, a box was drawn around each individual bound band and the apparent K_D_ was calculated for each individual binding event via a multi-state binding equation using the Solver function in Excel, which minimizes the error between the measured fraction bound values for each bound species and modeled fraction bound values based on a theoretical distribution. As an example of the calculated fraction bound in a gel with three bound species:

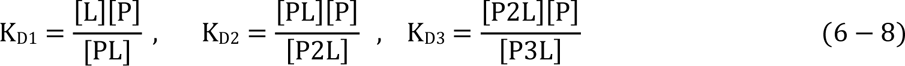

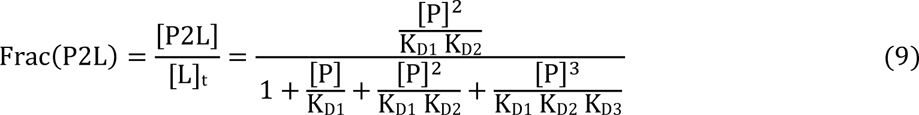

Where [L] is free RNA concentration, [P] is the concentration of unbound protein, [PL] is the concentration of the 1:1 protein:RNA complex, [P2L] is the concentration of the 2:1 protein:RNA complex, [P3L] is the concentration of the 3:1 protein:RNA complex, [L]_t_ is the total RNA concentration, and Frac(P2L) is the theoretical fraction of the 2:1 protein:RNA complex at a given protein concentration.

## RESULTS

### PureCLIP analysis of eCLIP data reveals two RNA motifs at hnRNP U binding sites

eCLIP datasets from the ENCODE consortium were analyzed to define a set of biologically relevant RNA targets of hnRNP U in cells as the basis for assessing the RNA sequence and/or structural features recognized by hnRNP U.^25^ Experiments performed in two cell lines, HepG2 and K562, with two biological replicates per cell line, were chosen to serve as the basis for defining hnRNP U-binding RNAs.^28^ Consistent with previous CLIP-based analyses of hnRNP U binding profiles,^8, 36^ the hnRNP U eCLIP reads were broadly dispersed across many transcripts, making it difficult to identify high-confidence binding sites through traditional peak callers. To help identify high-confidence hnRNP U direct interaction sites, we used PureCLIP, a peak caller that identifies protein-RNA crosslink sites based upon pileups of reverse-transcriptase read stops in eCLIP data.^26^ The criteria used by PureCLIP are more stringent than those used by other common peak callers, making it a necessary tool for identification of high-confidence binding sites within broadly dispersed eCLIP datasets.^37^ However, even after application of PureCLIP to identify hnRNP U crosslink sites in all four datasets, thousands of potential RNA targets were still identified, reinforcing prior findings of a broad distribution of protein-RNA interaction sites. To identify the highest-confidence binding sites, we filtered out any sites not observed across all four datasets and removed any sites with a signal:background ratio of <1, resulting in a final set of 323 crosslink sites (**Figure 2A**). Of the final 323 sites, 200 map to RNAs annotated by RefSeq,^38^ while the remaining 123 sites map to regions that do not match annotated RNAs.

**Figure 2.**
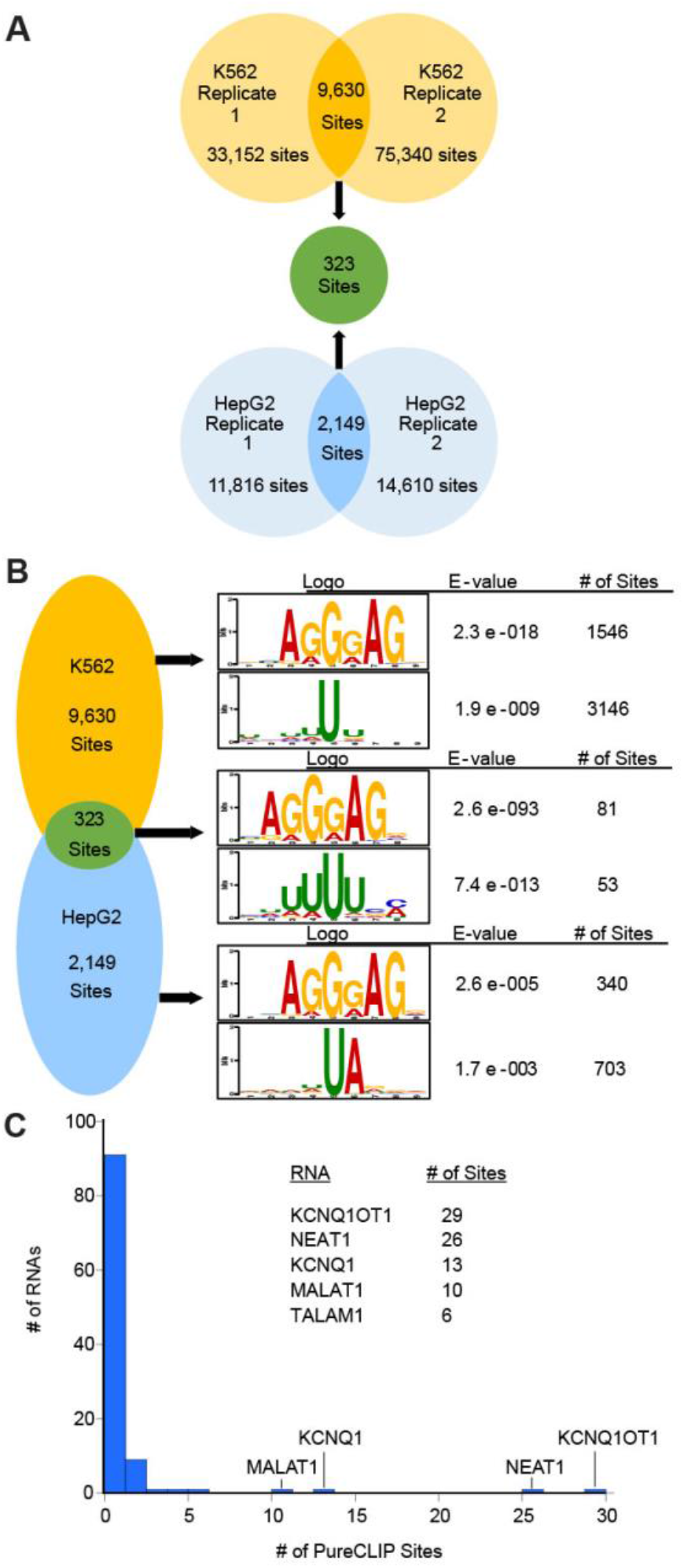
PureCLIP identifies RNA targets of hnRNP U. (A) Schematic representing the identification of hnRNP U crosslink sites conserved across all eCLIP biological replicates. (B) The AGGGAG and U-rich motifs are observed across both cell lines. Motifs within the final 323 sites were identified by MEME,^28^ while motifs within the larger K562 and HepG2 pools were identified by STREME.^29^ (C) Plot showing the number of PureCLIP sites within each annotated RNA containing at least one PureCLIP site.

The regions flanking each crosslink (+4 nucleotides in each direction) were searched for any enriched sequence motifs that may indicate preferred hnRNP U binding sites. MEME^28^ identified two main motifs enriched near the crosslink sites: 5’-AGGGAG and a uridine-rich (U-rich) motif (**Figure 2B**). Similar motifs were also enriched in the broader K562 and HepG2 datasets, indicating that sequence preferences were not lost in the process of filtering out lower-confidence binding sites. In the final list of 323 hnRNP U binding sites, 81 contained the 5’-AGGGAG motif, 53 contained the U-rich motif, and 189 did not contain either motif (**Figure 2B**). When the crosslink sites were further extended (+10 nucleotides in each direction), MEME identified the same two motifs, as well as a general G-rich motif with lower significance compared to the 5’-AGGGAG and U-rich motifs (**Figure S2**). The appearance of the G-rich motif within the extended crosslink sites may indicate that extended guanine-tracks broadly enhance hnRNP U occupancy at RNA sites.

### Selection of RNA ligands

After identifying sequence motifs enriched among RNA targets of hnRNP U, the set of 323 high-confidence crosslink sites was further narrowed to generate representative RNA targets for biochemical characterization. To accomplish this, RNAs were ranked by their PureCLIP score, which accounts for the number of reads ending at each nucleotide position, as well as enrichment over the background dataset.^26^ The top-ranked positions were dominated by a small number of RNAs known to be highly abundant in the nucleus, including MALAT1, NEAT1, and KCNQ1OT1 (**Figure 2C**).^39–41^ To ensure that the RNAs chosen for further analysis were not biased towards low-affinity interactions with highly abundant RNAs, only the top-scoring site was chosen from each of these RNAs. Lower-abundance RNAs were chosen to ensure that diverse representatives of each motif were selected. From the list of 323 RNA crosslink sites, a set of eleven RNA targets was selected (**Table 1**). This set of RNAs includes the observed 5’-AGGGAG and U-rich motifs, along with several RNAs that do not contain either motif.

**Table 1:**
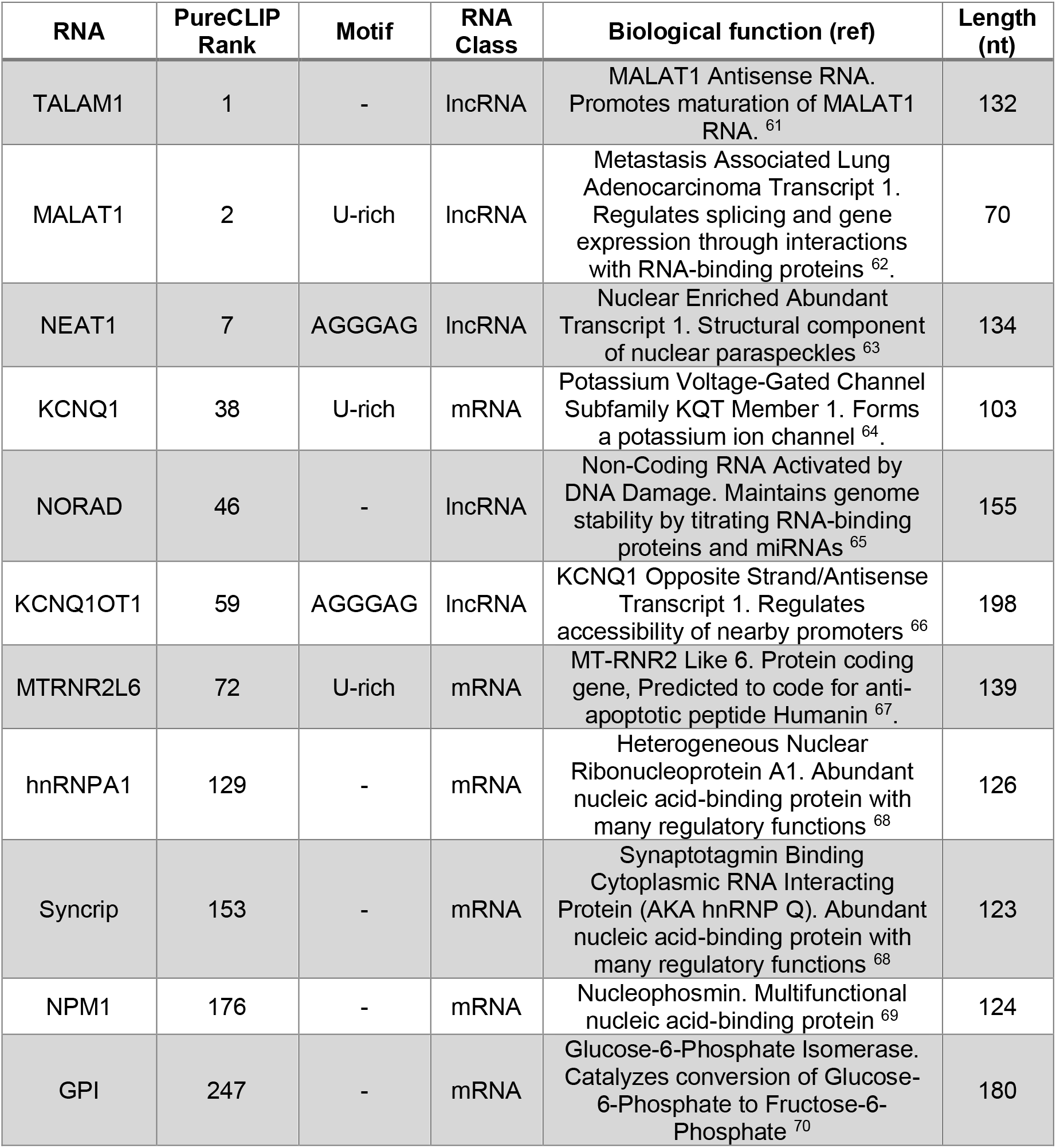
RNA fragments selected for biochemical characterization with hnRNP U.

To facilitate the design of the RNA fragments to be used for binding analysis, 200-nucleotide windows centered upon each selected crosslink site and the RNA structures of the resulting fragments were calculated using the Mfold algorithm accessed through the UNAfold web server.^31, 42^ These RNAs were further trimmed to encompass the regions of secondary structure around the crosslink site and their sequences were further refined in Mfold (**Figure 3A,B, Figure S3**).^31^ Analysis of the raw eCLIP alignments in the Integrative Genomics Viewer (IGV)^30^ confirmed that the minimized RNA fragments contained the full hnRNP U-bound regions proximal to the PureCLIP crosslink sites (**Figure 3C**). These eleven RNAs were synthesized by T7 RNAP and labeled at the 3’ end with FTSC for binding analysis.

**Figure 3.**
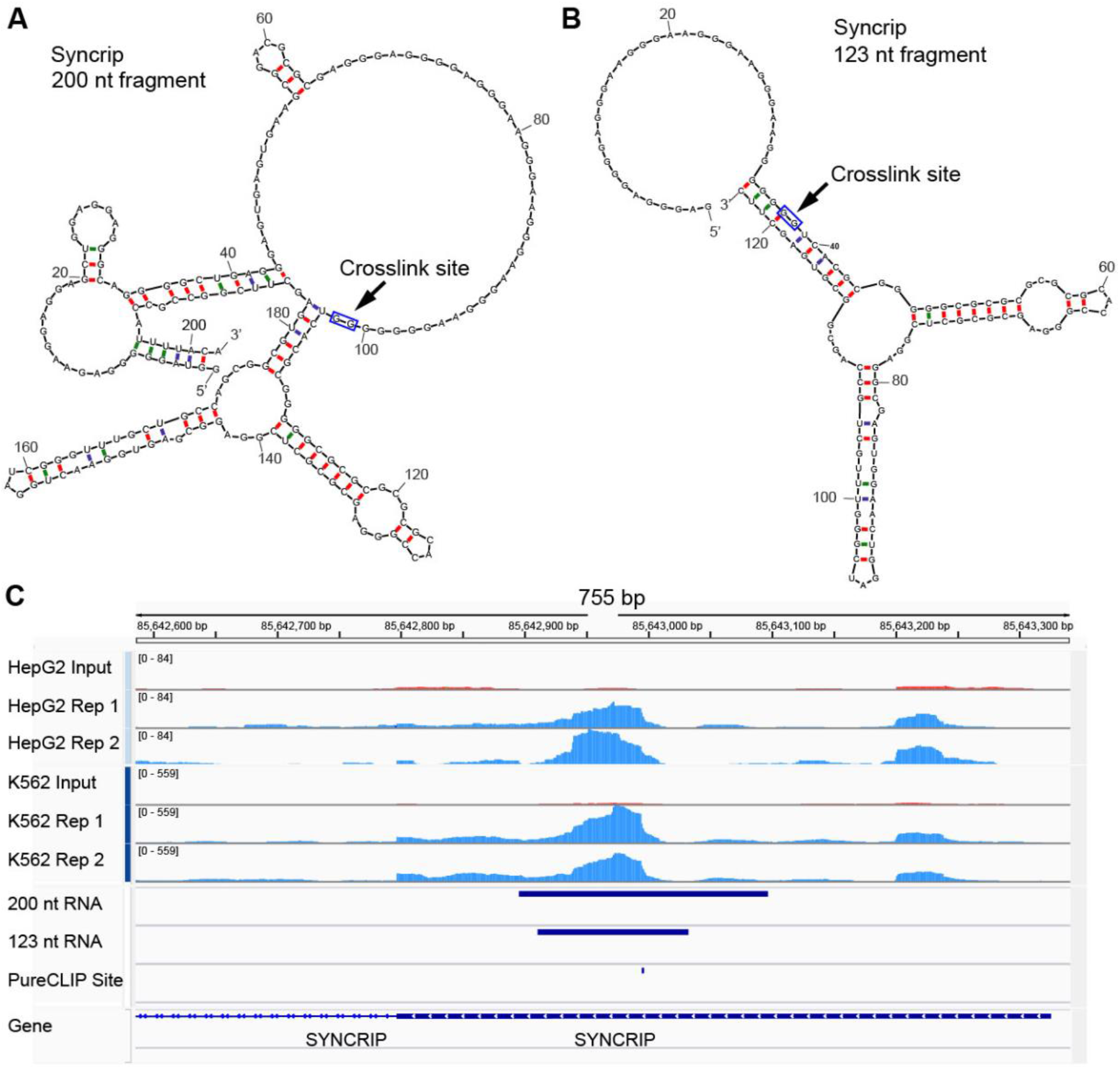
Design of minimized RNA fragments. (A) Initial Mfold structure prediction of the RNA from a 200 nt window centered around the Syncrip crosslink site. (B) Mfold structure prediction of the final 123 nt Syncrip fragment. (C) IGV traces showing raw eCLIP alignments alongside the Syncrip survey RNA. The low level red input traces were used as background for PureCLIP. The crosslink site identified by PureCLIP and the 200 and 123 nt fragments are shown in dark blue.

### hnRNP U binds RNA through its disordered C-terminus

To assess protein-RNA interactions, binding of FTSC-labeled RNAs to full length human hnRNP U with an N-terminal His_8_-MBP-tag were used in a fluorescence anisotropy (FA) equilibrium binding assay (note that throughout this study all proteins used for binding studies retained the His_8_-MBP tag). This assay was used because it enables high-throughput measurement of binding affinities in solution without disrupting the equilibrium between the bound and unbound species.^35, 43^ In this study, we do not make any initial assumptions from prior studies with regards to the RNA-binding elements within hnRNP U. Thus, we started our binding analysis with full length protein and minimized from there using a stepwise strategy. Full length hnRNP U bound all 11 RNAs with observed binding dissociation constants (K_obs_) spanning a ∼200-fold range from 11 – 2200 nM (**Table 2, Figure S4**). Note that while the observed binding for full length hnRNP U binding various RNAs could be fit with a binding equation that does not include the Hill coefficient (N_H_), for consistency of binding data analysis throughout this study all fitting was done with binding equations that contain the Hill coefficient to compare the behavior of different constructs. The Syncrip and MTRNLR2L6 RNAs were also tested for hnRNP U binding by electrophoretic mobility shift assays (EMSAs), which yielded similar values for K_obs_ (17±4 nM and 82±2 nM, respectively; **Figure S5**). These data indicate that hnRNP U can selectively bind RNA, but the motifs identified by MEME (5’-AGGGAG and U-rich) are not the only drivers of selective binding, as we observed no correlation between binding affinity and the presence of either motif. For further detailed analysis, we selected Syncrip and MTRNR2L6 as representative high- and mid-affinity RNAs, respectively.

**Table 2:**
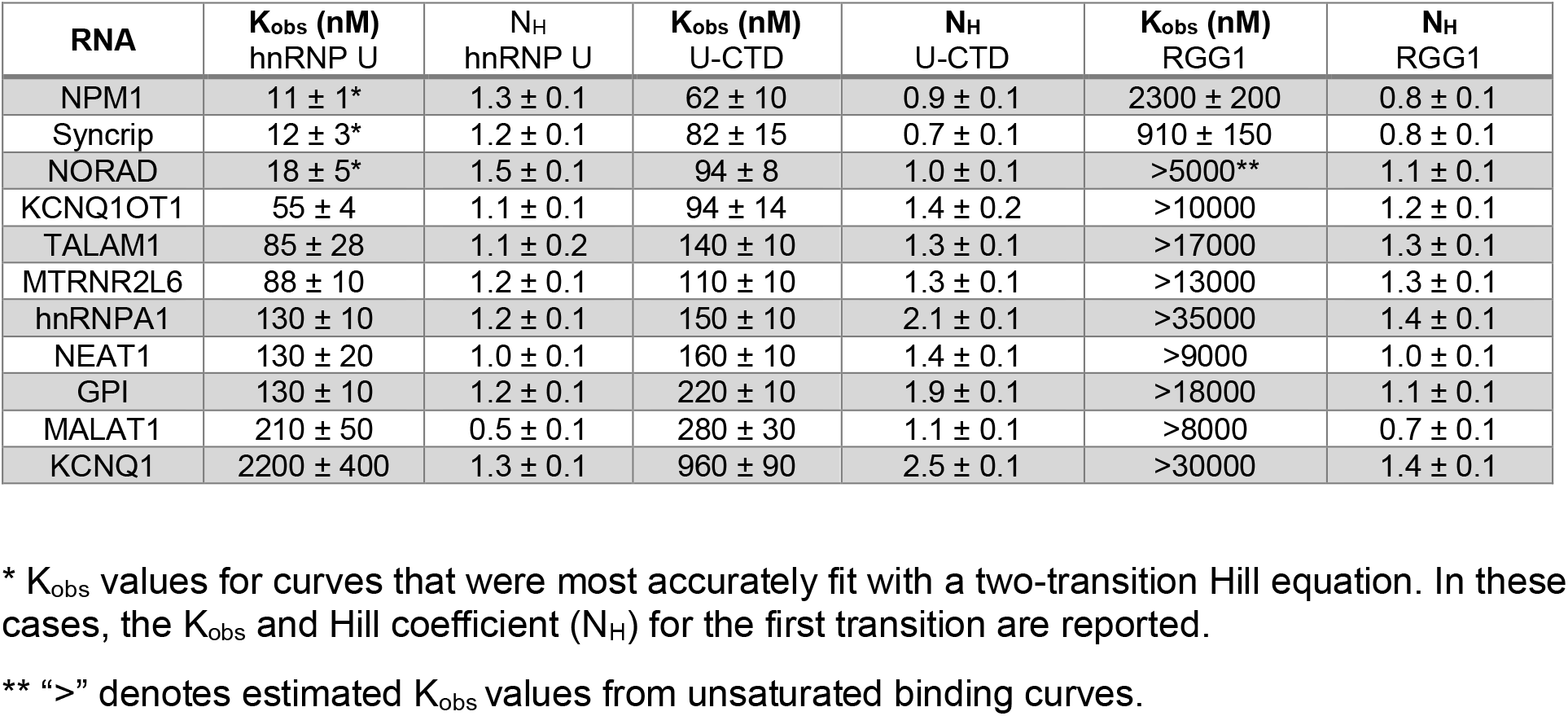
Apparent binding affinities of full-length hnRNP U, U-CTD, and RGG1.

Prior studies indicate that hnRNP U binds RNA through a disordered C-terminal region.^16^ A His_8_-MBP-tagged fragment corresponding to the complete C-terminal region of hnRNP U following the AAA+ domain (amino acids 673-825, referred to as U-CTD; **Figure 1**), which is predicted to be an IDR by both IUPRED2A and Alphafold^44–45^ (**Figure 4A, Figure S6**), was tested against the panel of 11 RNAs. The complete C-terminal region closely recapitulated the observed binding affinities and selectivity of the full-length protein (**Figure 4B**, **Table 2**), whereas the His_8_-MBP-tag alone did not bind any of the eleven RNAs, confirming that all observed binding is driven by the U-CTD (**Figure S7**). Most RNAs were bound with observed binding constants of <200 nM, whereas the KCNQ1 RNA exhibited much weaker affinity for both proteins (K_obs_ = 960 and 2200 nM for U-CTD and hnRNP U, respectively). An exception to the high correlation between observed affinities for hnRNP U and the U-CTD was the three RNAs that exhibited the highest affinity binding to hnRNP U (NPM1, Syncrip and NORAD fragments); each of these RNAs displayed a ∼5-fold lower affinity for U-CTD. However, these lower affinities are similar to the observed affinities of the U-CTD for the other RNAs (**Table 2**). Compared to the full-length protein, U-CTD exhibited a larger range of Hill coefficients (N_H_), from 0.7 to 2.5 (**Table 2**). However, for most RNAs, the Hill coefficient was between 1.0 and 1.5, comparable to the Hill coefficients exhibited by the full-length protein. It should be noted that the Hill coefficient, while often used as a measure of cooperative binding to describe highly defined molecular interactions, should not be interpreted as cooperative binding by hnRNP U or the RBD to RNA (see below). This parameter provides little insight into molecular mechanisms of binding equilibria^46^ and thus we consider it simply an empirical parameter in analyzing and interpreting our binding data.

**Figure 4.**
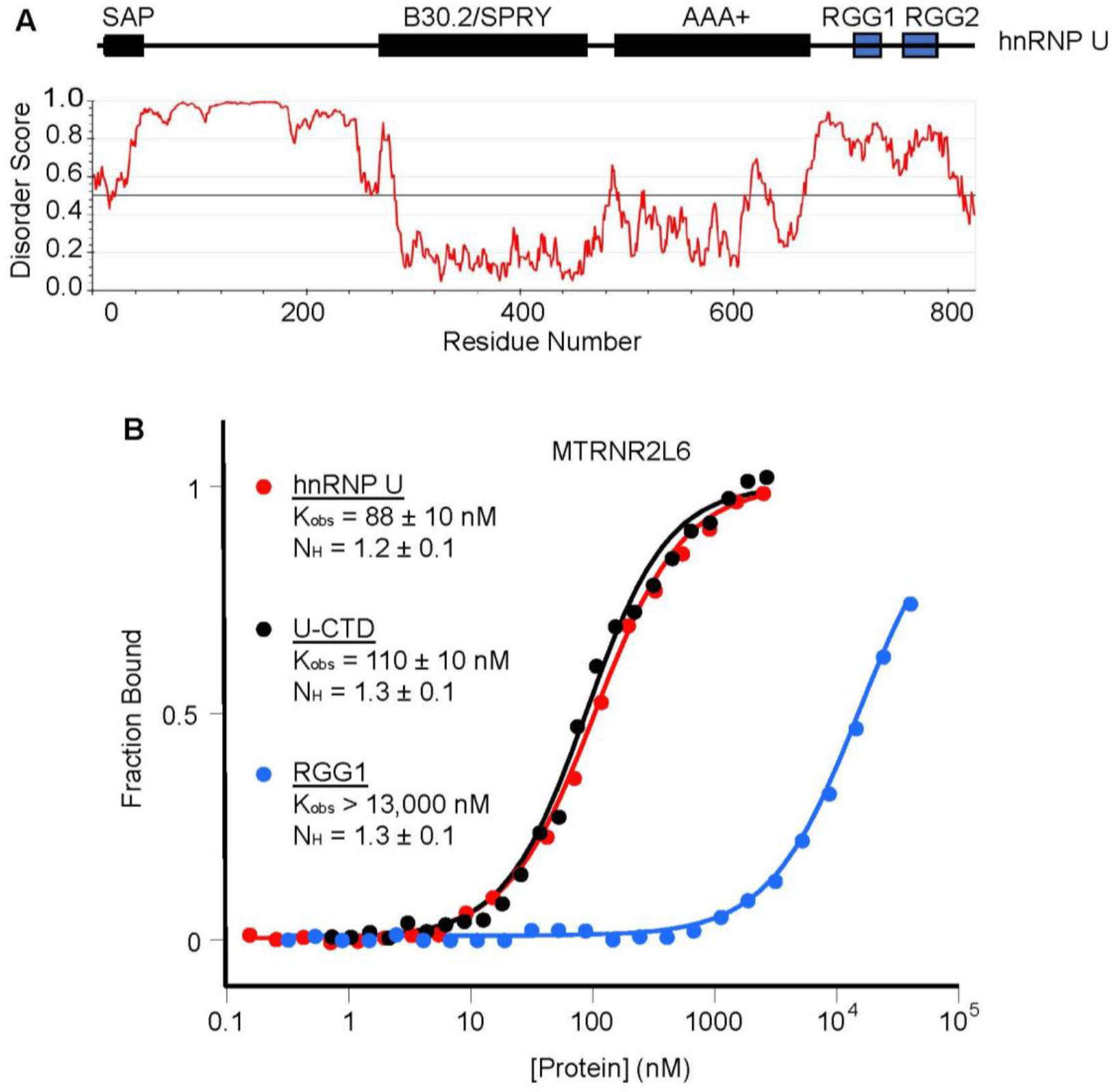
hnRNP U binds RNA through its disordered C-terminus. (A) Domain map of full-length hnRNP U (isoform 2), U(673-825) and U(702-745). (B) Predicted disorder is shown, as calculated by IUPRED2A.^44^ (C) Representative normalized FA curves of full-length hnRNP U, U(673-825) and U(702-745) bound to MTRNR2L6 RNA. Average K_obs_ values and Hill coefficients (N_H_) are shown with associated standard error of the measurement (s.e.m.).

Overall, our data correlates with previously reported measurements for hnRNP U-RNA interactions. These measurements include an observed 480 nM affinity for a dsRNA ligand measured by microscale thermophoresis,^47^ 2600 nM affinity for a 935-nt fragment of the *Xist* lncRNA measured by EMSA,^15^ 20-30 nM for a set of fragments of the *MALT1* mRNA^24^ and affinities ranging from 250 nM to >20 μM for a diverse panel of RNAs, as measured by EMSA.^9^ Although most of these studies did not report Hill coefficients, the interaction between a C-terminal fragment of hnRNP U and a fragment of *Xist* RNA exhibited a Hill coefficient of 1.1,^15^ comparable to our observed values. In contrast, another study measured Hill coefficients of 3-5 for fragments of *MALT1* pre-mRNA, indicative of cooperative, multisite binding by hnRNP U.^24^

### The canonical RGG1 motif is not the full hnRNP U RNA-binding domain

A previous biochemical study defined the RNA-binding domain of hnRNP U as a limited arginine- and glycine-rich region spanning amino acids 714-739, referred to as the “RGG box” (**Figure 1**) suggesting we might be able to further minimize beyond the U-CTD.^16^ However, this study analyzed binding of hnRNP U to simple RNA homopolymers and did not provide equilibrium binding affinity measurements, making it difficult to compare with binding affinity measurements provided by other studies. To determine if this region constitutes the full hnRNP U RNA-binding domain, a smaller protein fragment spanning this region along with several flanking amino acids (RGG1, **Figure 1**) was expressed and purified as the His_8_-MBP fusion and its RNA-binding properties assessed using the FA assay. We observed that the minimal RGG/RG element bound the panel of survey RNAs with significantly weaker affinity (11- to >120-fold change in affinity, **Table 2**) than the U-CTD, indicating that RGG1 in isolation does not recapitulate RNA binding by hnRNP U (**Figure 4B**, **Table 2**). These results reveal that the canonical RGG/RG element of hnRNP U is part of a larger RNA-binding domain, with amino acids important for recognition distributed throughout the disordered C-terminal domain.

### The U-CTD binds RNA through multivalent interactions

Because the FA measurements performed with the hnRNP U-CTD were best fit with a range of Hill coefficients, we questioned whether the variation in measured binding affinities between RNAs reflects multiple binding events. To visualize multi-site binding within hnRNP U-RNA interactions, EMSAs were performed with FTSC-labeled RNAs.^35^ Experiments with the His_8_-MBP-tagged U-CTD revealed that the protein-RNA complexes did not enter the gel, suggesting formation of higher-order complexes under these experimental conditions (**Figure S8A**). Because the very C-terminal end of hnRNP U is rich in aromatic amino acids and glutamines, which are known to promote protein-protein interactions,^48–49^ we reasoned that deletion of these region should enable intermediate RNA-protein complexes to enter the gel. Removal of the 16 C-terminal amino acids to generate the U-CTD(Δ16) fragment (**Figure 1, Figure S8B**) formed protein-RNA complexes that enabled visualization of several intermediate complexes on EMSAs in the 50-250 nM concentration regime, although higher order complexes still form above 500 nM concentration (**Figure S8C**). FA measurements with the panel of survey RNAs confirmed that this protein construct retains full RNA-binding activity, as most K_obs_ values were still within 2-fold of U-CTD (**Figure S8D**). Comparison of EMSAs of U-CTD and U-CTD(Δ16) suggest the C-terminal residues may promote protein-protein interactions along the RNA that yield cooperative binding behavior resulting in formation of higher-order protein-RNA complexes. In the absence of this sequence, individual RBDs tend to bind independently, resulting in the discrete intermediate complexes as observed by EMSA.

Using U-CTD(Δ16), EMSAs were performed with all eleven survey RNAs (**Figure 5A,B, Figure S9**). Multiple bound bands were observed with all RNAs, indicating that the hnRNP U RBD binds multiple sites within each RNA. Because these RNAs do not share a common structural motif, this suggests that the U-CTD interacts with a broad range of RNA sites while still maintaining sub-micromolar affinity. Multiple distinct bands were observed at lower protein concentrations, enabling K_D,app_ measurements of each binding event, although all RNA-protein complexes were too large to enter the gel when the protein concentration exceeded 1 μM. Some RNAs, such as Syncrip, appear to contain a single high-affinity binding site that saturates before weaker sites are bound (**Figure 5A**). Other RNAs, such as MTRNR2L6, appear to contain multiple equivalent binding sites, as evidenced by the fact that no individual site reaches saturation before multiple bands are observed (**Figure 5B**). Together, these EMSAs reveal that hnRNP U binds a broad range of RNA sites with high affinity with some RNAs apparently contain preferred binding sites, while others contain a set of near equal affinity sites. This is consistent with the general range of affinities observed in other studies.

**Figure 5.**
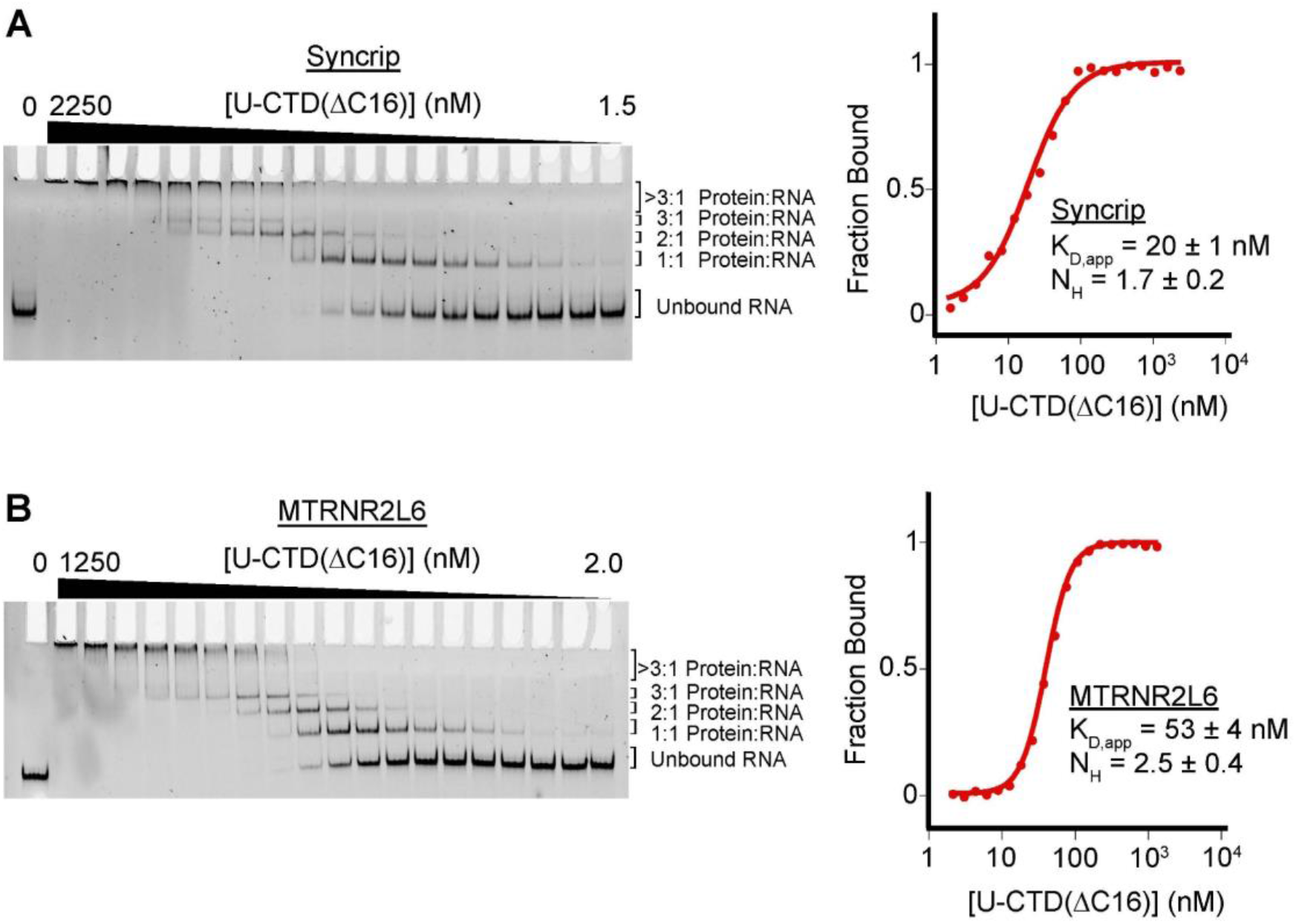
EMSA binding assays reveal intermediate protein-RNA complexes. EMSAs are shown alongside representative normalized binding curves with average K_D,macro_ and Hill coefficient. Examples are shown for (A) U-CTD(Δ16) bound to Syncrip RNA and (B) U-CTD(Δ16) bound to MTRNR2L6 RNA.

Two model RNAs, Syncrip and MTRNR2L6, were selected for more in-depth analysis of their binding stoichiometries. Using U-CTD(Δ16), EMSAs were performed in triplicate against both RNAs (**Figure 5A,B**), each individual band observed in the EMSA was quantified and the microscopic equilibrium dissociation constant (K_D,micro_) of each individual binding event was determined using a multi-state fitting function (**Table 3**). In addition, a single macroscopic dissociation constant (K_D,macro_) was calculated by considering all individual bound states as a single bound state. For Syncrip, the first transition represents a higher affinity interaction (∼20 nM each), while the second two are distinctly weaker (84 and 150 nM), suggesting non-equivalent binding sites within this RNA. In contrast, we observe nearly identical K_D,micro_ values for each binding event with MTRNR2L6, likely reflecting a set of equal and independent binding sites. These data reveal that hnRNP U, while binding most RNAs with mid-nanomolar affinity on the macroscopic scale, interact with them differently on the microscopic level, which may be the result of differences in how the two RGG/RG elements engage RNAs.

**Table 3:** EMSA affinity measurements.

| Protein<br>RNA | U-CTD($\Delta 16$) | |
| --- | --- | --- |
|  | Syncrip | MTRNR2L6 |
| $K_{D, \text{Micro-1}}$ (nM) | $22 \pm 2$ | $81 \pm 8$ |
| $K_{D, \text{Micro-2}}$ (nM) | $84 \pm 5$ | $88 \pm 4$ |
| $K_{D, \text{Micro-3}}$ (nM) | $150 \pm 40$ | $95 \pm 7$ |
| $K_{D, \text{Macro}}$ (nM) | $20 \pm 1$ | $53 \pm 4$ |
| $N_H$ | $1.7 \pm 0.2$ | $2.5 \pm 0.4$ |

### The hnRNP U RNA-binding domain contains a second RGG/RG element

To further examine potential differences between the two RGG/RG elements in the U-CTD, we first performed a binding analysis of the isolated RGG2 motif. A previous bioinformatic survey of human proteins for RGG/RG motifs predicted, but did not experimentally validate, a second potential RGG/RG element within the C-terminus of hnRNP U (RGG2), which most accurately fits the definition of a Tri-RG (more than two repeats of RG with 0-4 amino acids between each repeat) (**Figure 6A**).^17^ Comparison of the sequences of the two motifs shows that that RGG2 is almost entirely composed of RG repeats as opposed to RGG1, which is mostly composed of RGG repeats, and that RGG2 is more enriched in aromatic residues than RGG1.

**Figure 6.**
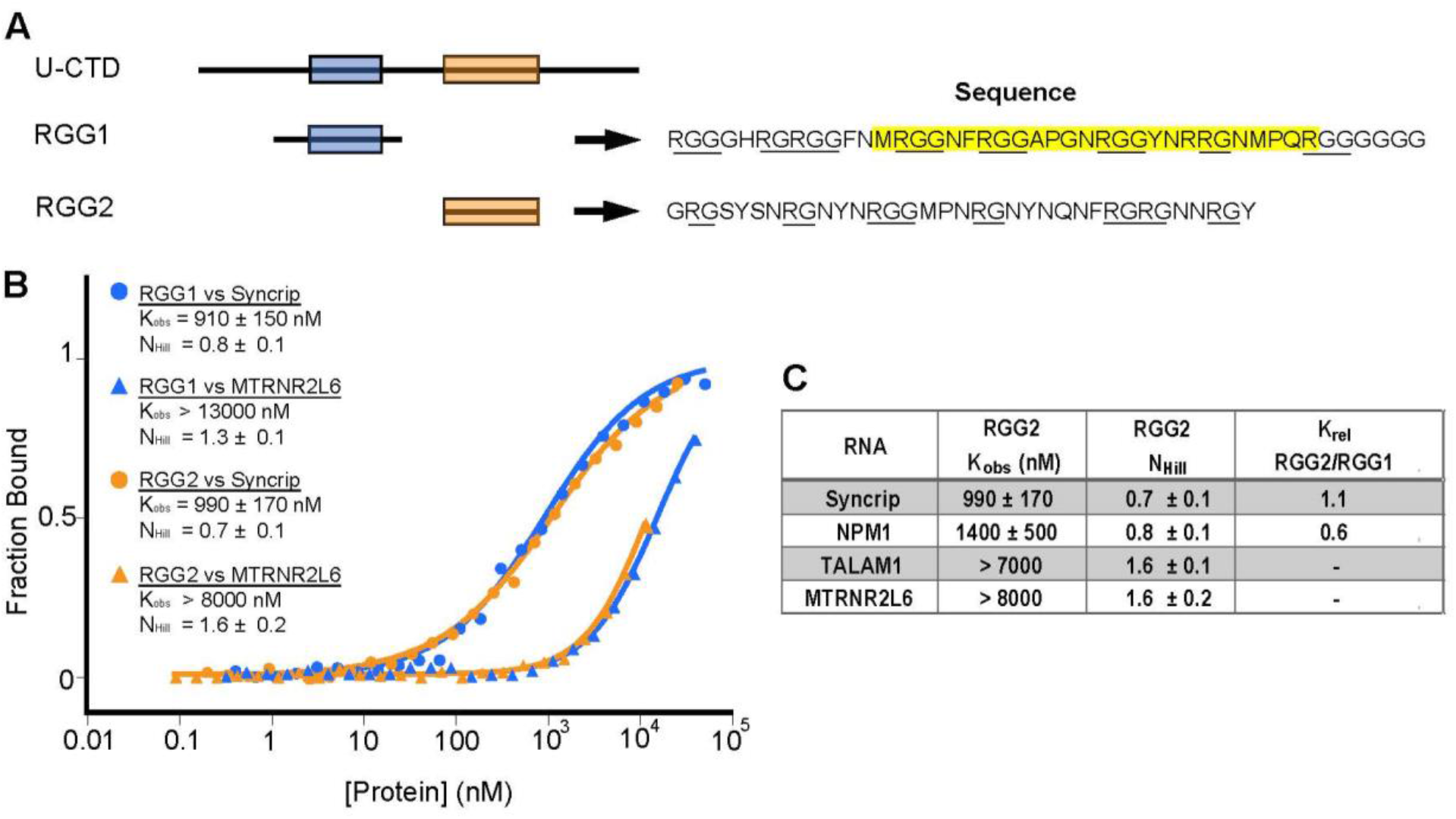
The hnRNP U RBD contains a second RGG/RG element. (A) Schematic of RGG1 and RGG2. The blue box represents the canonical RGG/RG element, and the cyan box represents the second RGG/RG element. Amino acid sequences of both fragments are included for reference. All RGG and RG repeats are underlined. The canonical RGG/RG element is highlighted in yellow. (B) Representative normalized FA curves of RGG1 or RGG2 bound to Syncrip or MTRNR2L6 RNAs, with average Hill coefficients and K_obs_. “>” denotes estimated K_obs_ values from unsaturated binding curves. (C) Table of mean K_obs_ and Hill coefficients for RGG2, with associated s.e.m. K_rel_ is a comparison with RGG1. “>” denotes estimated K_obs_ values from unsaturated binding curves.

To assess whether this region constitutes an authentic second RNA-binding RGG/RG element, a protein fragment corresponding to this region was expressed and purified. Equilibrium FA assays revealed that, despite the differences in amino acid composition between the two regions, this region binds RNA with similar binding affinities and RNA preferences as compared to the canonical RGG/RG element (RGG1) (**Figure 6B**). The K_obs_ for both RGG/RG elements binding to Syncrip and NPM1 are in close agreement with ITC measurements of the first RGG/RG element to hairpins derived from MALAT1 mRNA (0.6 – 1.1 µM).^24^ Together, these data confirm that the region of hnRNP U spanning amino acids 761-795 (Uniprot Q00839) contains a second functional RGG/RG element.

### RGG/RG elements serve redundant roles in RNA binding

To assess the individual contributions of RGG1 and RGG2 to RNA binding by the U-CTD, we generated a set of protein mutants that either deleted each RGG/RG element or mutated all arginine residues to serine (referred to as “SGG”) in the context of U-CTD while keeping the rest of the C-terminus intact (**Figure 7A**). We performed FA assays to compare binding preferences for a representative set of four RNAs (Syncrip, NPM1, MTRNR2L6, and TALAM1) from the panel of eleven survey RNAs. We found that removal of either individual RGG/RG element led to only modest losses of affinity for RNA (2.5- to 5-fold) (**Figure 7B,C**, **Table 4**). This indicates that a single RGG/RG element within the context of the CTD can confer most of the affinity for RNA in the absence of the other and that neither one is essential for RNA binding on its own. Thus, despite their sequence composition differences, RGG1 and RGG2 independently promote hnRNP U-RNA interactions. Further, it is important to note that the binding affinity for Δ-RGG1 is significantly greater than that of RGG2, and likewise for Δ-RGG2 versus RGG1. These data reveal that amino acid sequence flanking the RGG/RG elements in the CTG are important for establishing a high affinity interaction.

**Figure 7.**
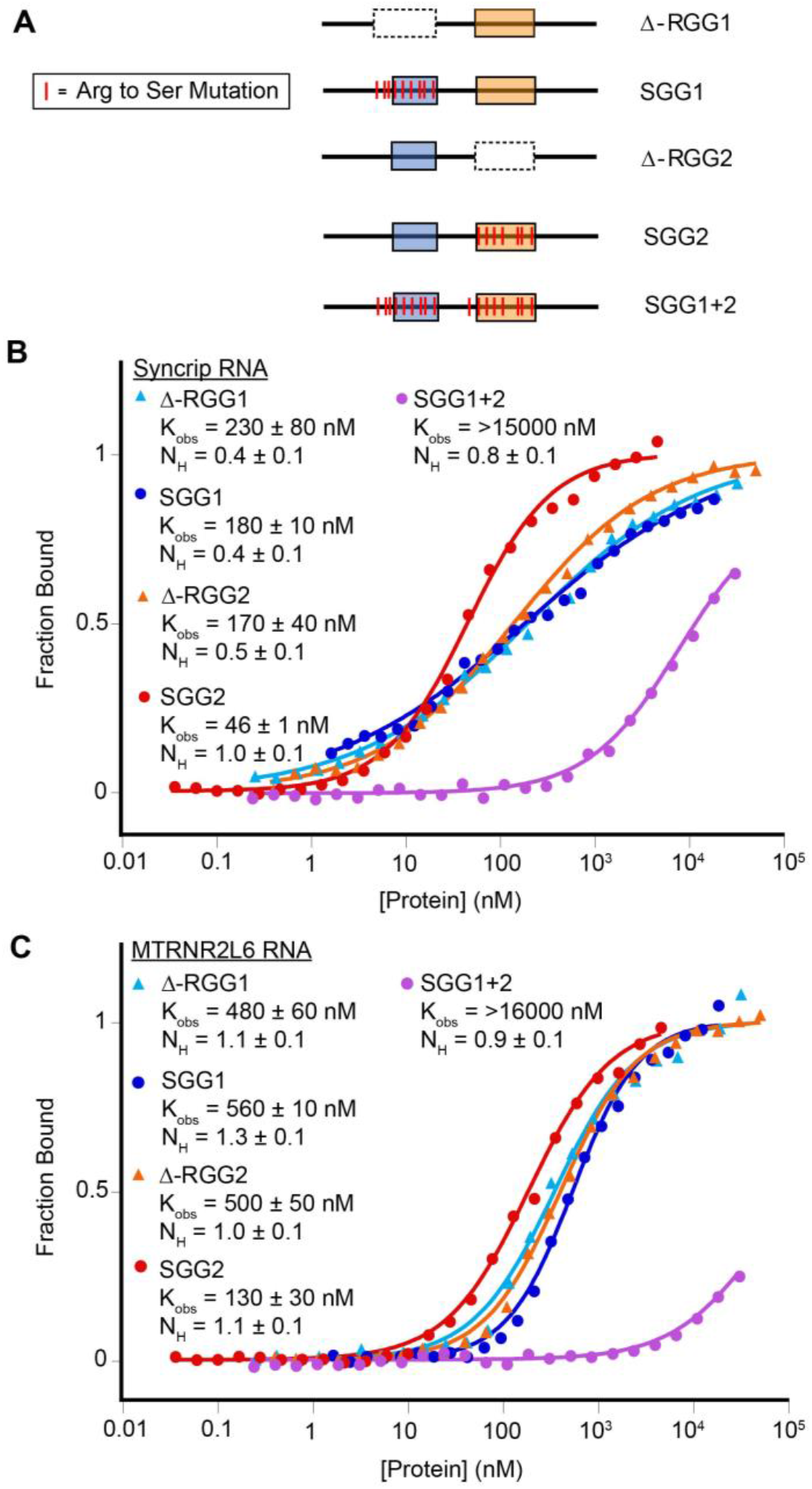
RGG/RG elements serve redundant roles in RNA binding. (A) Schematic of hnRNP U mutant and deletion constructs. Arg to Ser mutations are shown as red lines, and deleted regions are shown with a dotted line. (B) Representative normalized FA curves of hnRNP U mutant and deletion constructs bound to Syncrip RNA. Average K_obs_ values and Hill coefficients are shown with associated s.e.m. (C) Representative normalized FA curves of hnRNP U protein constructs bound to MTRNR2L6 RNA. Average K_obs_ values and Hill coefficients are shown with associated s.e.m.

**Table 4:** Apparent binding affinities of RGG/RG variants.

| RNA | Value | Δ-RGG1 | SGG1 | Δ-RGG2 | SGG2 | SGG1+2 |
| --- | --- | --- | --- | --- | --- | --- |
| **NPM1** | $K_{obs}$ | $160 \pm 20$ | $210 \pm 10$ | $230 \pm 40$ | $53 \pm 7$ | $>13000^{**}$ |
| | $K_{rel}^*$ | 2.6 | 3.6 | 3.7 | 0.9 | $>200$ |
| | $N_H$ | $0.5 \pm 0.1$ | $0.5 \pm 0.1$ | $0.4 \pm 0.1$ | $0.8 \pm 0.1$ | $0.8 \pm 0.1$ |
| **Syncrip** | $K_{obs}$ | $230 \pm 80$ | $180 \pm 10$ | $170 \pm 40$ | $46 \pm 1$ | $>15000$ |
| | $K_{rel}$ | 2.9 | 2.3 | 2.1 | 0.6 | $>180$ |
| | $N_H$ | $0.4 \pm 0.1$ | $0.4 \pm 0.1$ | $0.5 \pm 0.1$ | $1.0 \pm 0.1$ | $0.8 \pm 0.1$ |
| **MTRNR2L6** | $K_{obs}$ | $480 \pm 60$ | $560 \pm 10$ | $500 \pm 50$ | $130 \pm 30$ | $>16000$ |
| | $K_{rel}$ | 4.4 | 5.0 | 4.5 | 1.2 | --- |
| | $N_H$ | $1.1 \pm 0.1$ | $1.3 \pm 0.1$ | $1.0 \pm 0.1$ | $1.1 \pm 0.1$ | $0.9 \pm 0.1$ |
| **TALAM1** | $K_{obs}$ | $530 \pm 50$ | $610 \pm 10$ | $500 \pm 70$ | $140 \pm 10$ | N.D.B.*** |
| | $K_{rel}$ | 3.8 | 4.4 | 3.6 | 1.0 | N/A |
| | $N_H$ | $1.1 \pm 0.1$ | $1.5 \pm 0.1$ | $1.2 \pm 0.1$ | $1.1 \pm 0.1$ | $0.7 \pm 0.1$ |

\* $K_{rel}$ denotes a comparison with $K_{obs}$ measured for U-CTD
\*\* ">" denotes estimated $K_{obs}$ values from unsaturated binding curves.
\*\*\* N.D.B.: No Detectable Binding

The role of arginine residues in the two RGG/RG motifs was examined using a set of arginine-to-serine mutations (**Figure 7A**). The SGG1 mutant had a similar effect as the deletion of the first RGG/RG element, indicating that arginines substantially drive RNA recognition by the RGG1. In contrast, the SGG2 mutant had no discernable effect on affinity in contrast to deletion of RGG2 which led to modest losses of affinity (2- to 4.5-fold loss in affinity). This indicates that other amino acids play a larger role in RNA recognition by RGG2 (**Figure 7B,C**), indicating that RNA recognition by RGG/RG elements can differ, suggesting a role of amino acids beyond arginine in mediate RNA contacts. However, underscoring the overall importance of arginine residues in mediating hnRNP U’s interactions with RNA, mutation of all arginine residues through the C-terminus (SGG1+2) induced a severe to near-complete loss of affinity for RNA (**Figure 7B,C**).

To directly visualize formation of discrete protein-RNA complexes by the above panel of mutants, EMSAs were performed against Syncrip and MTRNR2L6 RNAs. Importantly, EMSAs confirm that perturbing a single RGG/RG element does not substantially decrease affinity for RNA, as SGG1, SGG2, ΔRGG1 and ΔRGG2 bind Syncrip and MTRNR2L6 with K_D,app_ values below 250 nM (**Figure S10A,B**). These data reinforce that observation by FA that the two RGG/RG elements are at least partially redundant in that both ΔRGG mutants retain the ability to form discrete intermediate complexes, particularly with Syncrip RNA. Further, although no single set of arginine residues in this region is essential for RNA binding, mutation of all arginine residues to serine dramatically impairs the RBD’s ability to form stable RNA-protein complexes. Interestingly, a few of these protein constructs behave differently from U-CTD(Δ16) with respect to formation of discrete intermediate protein-RNA complexes observed by EMSA (left, **Figure S10C,D**). The SGG1 and individual RGG/RG deletion mutants bound to Syncrip RNA exhibit distinct intermediate complexes similar to U-CTD(Δ16). In contrast, the SGG2 and SGG1+2 mutants either form intermediate smeared bands (right, Figure **S10C,D**) or higher-order complexes that do not enter the gel, suggesting that these proteins behave differently with RNA due to the loss of arginine residues. For example, a change in their kinetic behavior can lead to the observed pattern of bands in gels due to dissociation of the complex during electrophoresis.^50–51^

## Discussion

hnRNP U is a highly abundant RNA-binding protein that serves many biological functions through its interactions with RNA.^3–4, 6^ While the importance of hnRNP U as a key player in the nuclear scaffold that links DNA and RNA together is well established, the molecular details of its ability to interact with numerous nuclear RNAs remains largely unexplored. Towards a detailed characterization of how hnRNP U binds RNA, we first determined the precise RNA-binding domain within this protein, revealing a set of key features of this RBD.

### The minimal RNA binding domain spans the C-terminal disordered region of hnRNP U

To experimentally define the RBD of hnRNP U, we established a set of biologically relevant RNA targets of hnRNP U using PureCLIP analysis of ENCODE eCLIP data, resulting in a panel of 11 RNAs representative of *in vivo* targets. Deletion analysis of hnRNP U with this panel of RNAs revealed that the necessary and sufficient protein domain that was able to maintain high affinity binding spans residues 673-809 (**Figure 1**). While removal of the last 16 amino acids of hnRNP U did not cause a significant loss in affinity, this deletion resulted in a distinct change in the binding behavior of the hnRNP U RBD as observed by EMSA. The full RNA binding domain, U-CTD, displayed binding behavior in which the RNA becomes saturated over a narrow protein concentration range, as evidenced by lack of intermediate protein-RNA complexes in favor of low mobility, higher-order complexes. Loss of the 16 C-terminal residues yields an RBD which forms discrete intermediate species prior to formation of the higher order complexes, suggesting that this region promotes protein-protein interactions and cooperative binding behavior. Importantly, further minimization of the U-CTD to the individual RGG/RG motifs (RGG1 and RGG2), severely reduced RNA binding activity, indicating that residues outside of these motifs play important roles in supporting high affinity binding.

### The hnRNP U RBD contains two distinct functional RGG/RG motifs

Because a recent bioinformatic survey identified a second potential RGG/RG element in the disordered C-terminal tail of hnRNP U,^17^ we examined the ability of the two elements to bind RNA. Comparison of RNA binding by each of the individual RGG/RG elements confirmed that the second hnRNP U RGG/RG element (RGG2) binds RNA with similar affinity to the first; RGG-2 displays approximately two-fold weaker affinity for RNA but shows a similar ∼4-fold preference for Syncrip over MTRNR2L6, as measured by EMSA. The similar affinities of these two RGG/RG elements are despite sequence differences between them; RGG2 is classified as a Tri-RG whereas RGG1 is identified as a Tri-RGG.^17^ Thus, the distinction between RGG- and RG-repeats do not appear to result in functional differences in their affinity for RNA, although the set of protein-RNA contacts used to achieve binding by RGG/RG elements may be different (see below).

### The two RGG/RG elements are functionally redundant

The observation that hnRNP U RBD contains two functional RGG/RG elements is consistent with a bioinformatic analysis demonstrating that most RGG/RG proteins contain multiple RBDs, with the RGG/RG typically adjacent to a structured RBD such as an RRM or KH domain.^18^ Deletion of each RGG/RG element in the context of the full C-terminal RBD demonstrated that hnRNP U can tolerate the loss of either RGG/RG element with only slightly reduced binding affinity. This suggests that both RGG/RG elements are functionally redundant with respect to direct RNA binding in the context of the full protein and is consistent with previous reports that RGG/RG elements act as affinity anchors that work in concert with nearby RBDs.^22, 52^ In the case of hnRNP U, the two RGG/RG elements support each other with neither serving as a “core” RBD, thereby acting as a bivalent RNA ligand. Although both RGG/RG elements work together, it should be noted that the binding contributions of each RGG/RG element are not additive. The fact that both RGG/RG elements work together to moderately enhance binding affinity but do not appear to work in a purely additive manner points to influences upon the avidity effect such as spacing between binding sites within a particular RNA and the nature of the linker.^53–54^ Thus, our data suggests that hnRNP U uses two tandem RGG/RG elements with modest affinity for RNA to achieve high affinity binding, likely by enabling longer residence times by keeping the RNA-protein complex intact after one RGG/RG element dissociates^54^. This is consistent with the idea of RGG/RG elements acting as “affinity anchors” that confer affinity for RNA without strong bias for any particular targets.^52^ The unique use of tandem RGG/RG domains by hnRNP U without the assistance of adjacent structured RBDs is likely the critical feature that imparts its ability to broadly bind most nuclear RNAs to generate a stable nuclear scaffold.

### Arginine residues are the primary but not sole drivers of RNA binding

Previous investigations into the RNA-binding characteristics of RGG/RG elements have focused upon arginine residues as key drivers of affinity.^9, 52, 55^ Arginines in both RGG/RG elements and arginine rich domains of viral proteins can make both base-specific contacts through guanine and electrostatic contacts with the phosphate backbone of RNA.^56^ In the case of hnRNP U, complete arginine to serine mutation (R-to-S) of either RGG/RG element is only modestly detrimental binding, revealing the redundant nature of the RGG/RG domains. Mutating arginines throughout the C-terminal region induced a larger loss of affinity, but the SGG1+2 mutant RBD was still able to bind RNA with sub-micromolar affinity. This suggests that while arginine residues are primary drivers of RNA binding, the RBD contains other RNA-interacting residues. This is also consistent with the observation that the minimal peptides do not have the same RNA binding affinity as deletion of their counterpart in the context of the U-CTD, further supporting the role of non-arginine residues for RNA recognition.

The hnRNP U RBD is significantly enriched in amino acid residues that have a strong propensity to interact with nucleic acids. A detailed bioinformatic analysis of RNA-protein interfaces revealed that in addition to a high frequency of arginine, a set of other residues also have a relatively high occurrence including aromatic (Y and F) and polar (N and Q) residues.^57^ These residues also are found preferentially mediating protein-DNA interactions as well, indicating their general preference for interactions with nucleic acids.^58–59^ The composition of the U(673-809) region of hnRNP U, which comprises the minimal RNA binding domain, is substantially enriched in these residues along with arginine and glycine (R, 12.4%; G, 37%; N, 21%; Q, 7.2%; Y, 5.8%; F, 3.6%). The observation that the SGG1+2 mutants retain some RNA binding activity suggests these residues may play accessory roles in binding. These residues may serve to greatly expand the potential RNA binding sites of hnRNP U through base-specific and pi-stacking interactions not accessible by arginine residues. Thus, the RGG/RG domain may serve to scaffold a broad spectrum of general nucleic acid interacting residues within the context of an intrinsically disordered domain that gives it the flexibility to broadly recognize RNA.

Previous literature supports this idea; one study found that truncating the C-terminal tail induced mislocalization of the *Xist* lncRNA^4^ while a separate study reported no change in *Xist* localization upon precisely deleting the canonical RGG/RG element but not the rest of the C-terminal tail,^60^ which can be explained by residual RNA-binding activity by the rest of the RBD. Multi-RBD binding has been observed in the RGG/RG elements of FUS, which achieve strong affinity by working in tandem with each other and neighboring RRM domains.^9, 22^ In a similar vein, we found that the canonical RGG/RG element of hnRNP U works in tandem with a neighboring RGG/RG element to achieve high affinity for its RNA targets.

## ASSOCIATED CONTENT

### Supporting Information

The Supporting Information is available free of charge at (web address). Supporting Information contains materials and methods, supplementary Figures and Tables.

## Data availability

Data will be made available on request.

## Accession codes

HNRNPU: Q00839

## Author Contributions

O.A.K, D.S.W., and R.T.B. designed the research. O.A.K. conducted the experimental research and wrote the paper with feedback from all authors. D.S.W. and R.T.B. supervised the project.

## Funding

This work was supported by a grant from the National Institutes of Health to D.S.W. and R.T.B. (R01 GM120347) and a grant from the National Institutes of Health to R.T.B. (R01 GM073850) and by an National Institutes of Health T32 training grant (GM008759) through the University of Colorado Boulder.

## Notes

R.T.B. serves on the Scientific Advisory Boards of Expansion Therapeutics, SomaLogic and MeiraGTx.

## ACKNOWLEDGEMENTS

The authors thank the Shared Instrument Pool (SIP) core facility (RRID SCR_018986), Department of Biochemistry, University of Colorado Boulder, for the use of the Typhoon FLA 9500 and centrifuges. We would like to thank Dr. Annette Erbse for help designing the EMSA and FA assays.

## ABBREVIATIONS

EMSA: electrophoretic mobility shift assay
eCLIP: enhanced UV crosslinking and immunopurification
FA: fluorescence anisotropy
hnRNP U: Heterogeneous nuclear ribonucleoprotein U
IDR: intrinsically disordered region
RBD: RNA-binding domain

