## Supplemental Information for "The RNA-binding domain of hnRNP U extends beyond the RGG/RG motifs"

### Table of Contents

**Table S1:** Sequences of RNAs used in this study

**Table S2:** Sequences of proteins used in this study

**Figure S1:** SDS-PAGE analysis of protein purification

**Figure S2:** MEME motifs of the extended hnRNP U PureCLIP sites

**Figure S3:** Predicted secondary structures of survey RNAs

**Figure S4:** Representative FA curves of hnRNP U bound to survey RNAs

**Figure S5:** Representative EMSAs of hnRNP U bound to survey RNAs

**Figure S6:** Alphafold structure prediction of hnRNP U

**Figure S7:** MBP does not bind survey RNAs

**Figure S8:** Truncation of the C-terminus reduces well-shifting of protein-RNA complexes

**Figure S9:** EMSAs of survey RNAs

**Figure S10:** EMSAs reveal altered binding modes of RGG/RG mutants

**Supplementary Table 1:** Sequences of RNAs used in this study

| RNA | Length (nt) | Sequence |
|---|---|---|
| Syncrip | 123 | GAGGGAGGGGAGGGAAGGGAAGGGAAGGGAAGGGGGGUCACGCGGG<br>GGCGCGCGCGCGCACCGGGAGCGCGCUCGGAGGCGAGUGGAACUGGA<br>UCGGGUUUGCUGCCAGCGGCGUGAGCUUC |
| NPM1 | 124 | GGGUGGGGAGGCGCGGCGCGGCGUGAGGAGGCCGCAACCGCUGGGAG<br>CACGUGGUUGCCACGUGGUUGGGGGAGGGAGGGGGGUGUGGGCGCUA<br>CAUCCGGGACUCACCGGCGUGUUGCCACUC |
| NORAD | 156 | GAGACUAUUCAUUGGGUGUUUGGGGUGGGGGAAGGGGGGGUGGGCAG<br>AGGAGGUAUGCAGGGAGAGGGGUUCUGUGCUCCUGAGAUUAGUUCAG<br>AUGGUCUAACCAUUGUUCUAUAUGUGCAUUUUAGUUAAUAUUGUGUA<br>UUAAAGGAUAAGUCU |
| KCNQ1OT1 | 198 | GGAAGCAACGGCAGCCUGGUCUUUAAGGGUGAGGGAGCACCUUCUCC<br>UGAGAGAGGGAGAGAUAGGUUGGGGGAGGUUGGGUGGGGGUGGGGAG<br>AGCGAGUGAGCAAGCAAGCUAGAAACUUCCUGAAAAGGGAAGUUACU<br>CUAGAAAAGGGUAAGUUCCUUCCAGGCAUCAUGGGCUGGCUUAACCU<br>GGACCAGUUU |
| MTRNR2L6 | 139 | GCUAUACAUAUAUUGACCCAAUAAUUUGAUCAACGGAACAAGUUACC<br>CUAGGGAUAACAGCGCAAUCCUAUUCUAGAGUCCAUAUUGACAAUAG<br>GGUUUACGACCUCGAUGUUGGAUCAGGACAUCCUAAUGGUGUAGC |
| TALAM1 | 132 | GGAUGAAAUGAAGCAACUCUUCUGAUAACGAAGAGAUACCUGUCUGA<br>GGCAAACGAAACAUUGGCACACAGCACAGCCUCCUCAAUCCACUUGA<br>UCCCAACUCAUCUCUCAUUUAUUUCGGCUUCUUUUAUC |
| hnRNPA1 | 126 | GGUUUUAAUGUAGAUUUUUUUUUUUGCACCCCAUGCUGUUGAUUGCU<br>AAAUGUAACAGUCUGAUCGUGACGCUGAAUAAAUGUCUUUUUUUUAA<br>UGUGCUGUGUAAAGUUAGUCUACUCUUAAGCC |
| NEAT1 | 134 | GGCAGGGAGAGGUAGAAGGGGUGGAGGAGUCAGGAGGAAUAGGCCGC<br>AGCAGCCCUGGAAAUGAUCAGGAAGGCAGGCAGUGGGUGCAGGGCUG<br>CAGGAGGGCCGGGAGGGCUAAUCUUCAACUUGUCCAUGCC |
| GPI | 180 | GGGAAUGAGGAGAUGUGUCUUCCAGUGACUUAGGAGGUACGAUCUCU<br>UAUAGGUAUGAGGAUCUGGGUCUGUCAAUGUCUGAGGAGGUAGGAUC<br>UGGCCCUGGUAUGAGAAUCCGGGUCUGUCCAUGUCUGAGCAGGUAGG<br>AUGUGUCCCUGGUAUGAGGAUCCGGGUCUGUCAAUUUCC |
| MALAT1 | 70 | GUUGGCGUGGGGGUGGAGGGGUGAGGUGGGCGCUAAGCCUUUUUUUA<br>AGAUUUUUCAGGUACCCCUCACC |
| KCNQ1 | 103 | GGCCCUCAGUCUACACUCUGUGAUCCACUCAUGAAUAUCUGGUCUGG<br>CUCUGCGUUCCAGUCAGACCCUUUUCUGAGAGGACCACAAGUCAUAG<br>CCCAGGGCC |

**Supplementary Table 2:** Sequences of protein constructs used in this study

| Protein Construct | Sequence | Extinction Coefficient $M^{-1}cm^{-}$ | pI |
|---|---|---|---|
| hnRNP U | MHHHHHHHHKIEEGKLVIWINGDKGYNGLAEVGKKFEKDT<br>GIKVTVEHPDKLEEKFPQVAATGDGPDIIFWAHDRFGGYA<br>QSGLLAEITPDKAFQDKLYPFTWDAVRYNGKLIAYPIAVE<br>ALSLIYNKDLLPNPPKTWEEIPALDKELKAKGKSALMFNL<br>QEPYFTWPLIAADGGYAFKYENGKYDIKDVGVDNAGAKAG<br>LTFLVDLIKNKHMNADTDYSIAEAAFNKGETAMTINGPWA<br>WSNIDTSKVNYGVTVLPTFKGQPSKPFVGVLSAGINAASP<br>NKELAKEFLENYLLTDEGLEAVNKDKPLGAVALKSYEEEL<br>AKDPRIAATMENAQKGEIMPNIPQMSAFWYAVRTAVINAA<br>SGRQTVDEALKDAQTNSSSLVPRGSMSSSPVNVKKLKVSE<br>LKEELKKRRLSDKGLKAELMERLQAALDDEEAGGRPAMEP<br>GNGSLDLGGDSAGRSGAGLEQEAAAGGDEEEEEEEEEEEG<br>ISALDGDQMELGEENGAAGAADSGPMEEEEAASEDENGDD<br>QGFQEGEDELGDEEEGAGDENGHGEQQPQPPATQQQQPQQ<br>QRGAAKEAAGKSSGPTSLFAVTVAPPGARQGQQQAGGKKK<br>AEGGGGGGRPGAPAAGDGKTEQKGGDKKRGVKRPREDHGR<br>GYFEYIEENKYSRAKSPQPPVEEEDEHFDDTVVCLDTYNC<br>DLHFKISRDRLSASSLTMESFAFLWAGGRASYGVSKGKVC<br>FEMKVTEKIPVRHLYTKDIDIHEVRIGWSLTTSGMLLGEE<br>EFSYGYSLKGIKTCNCETEDYGEKFDENDVITCFANFESD<br>EVELSYAKNGQDLGVAFKISKEVLAGRPLFPHVLCHNCAV<br>EFNFGQKEKPYFPIPEEYTFIQNVPLEDRVRGPKGPEEKK<br>DCEVVMMIGLPGAGKTTWVTKHAAENPGKYNILGTNTIMD<br>KMMVAGFKKQMADTGKLNTLLQRAPQCLGKFIEIAARKKR<br>NFILDQTNVSAAAQRRKMCLFAGFQRKAVVVCPKDEDYKQ<br>RTQKKAEVEGKDLPEHAVLKMKGNFTLPEVAECFDEITYV<br>ELQKEEAQKLLEQYKEESKKALPPEKKQNTGSKKSNKNKS<br>GKNQFNRGGGHRGRGGFNMRGGNFRGGAPGNRGGYNRRGN<br>MPQRGGGGGGSGGIGYPYPRAPVFPGRGSYSNRGNYNRGG<br>MPNRGNYNQNFRGRGNNRGYKNQSQGYNQWQQGQFWGQKP<br>WSQHYHQGYY | 139,580 | 5.6 |
| U-CTD | MHHHHHHHHKIEEGKLVIWINGDKGYNGLAEVGKKFEKDT<br>GIKVTVEHPDKLEEKFPQVAATGDGPDIIFWAHDRFGGYA<br>QSGLLAEITPDKAFQDKLYPFTWDAVRYNGKLIAYPIAVE<br>ALSLIYNKDLLPNPPKTWEEIPALDKELKAKGKSALMFNL<br>QEPYFTWPLIAADGGYAFKYENGKYDIKDVGVDNAGAKAG<br>LTFLVDLIKNKHMNADTDYSIAEAAFNKGETAMTINGPWA<br>WSNIDTSKVNYGVTVLPTFKGQPSKPFVGVLSAGINAASP<br>NKELAKEFLENYLLTDEGLEAVNKDKPLGAVALKSYEEEL<br>AKDPRIAATMENAQKGEIMPNIPQMSAFWYAVRTAVINAA<br>SGRQTVDEALKDAQTNSSSVPGRGSSKKALPPEKKQNTGS<br>KKSNKNKSGKNQFNRGGGHRGRGGFNMRGGNFRGGAPGNR<br>GGYNRRGNMPQRGGGGGGSGGIGYPYPRAPVFPGRGSYSN<br>RGNYNRGGMPNRGNYNQNFRGRGNNRGYKNQSQGYNQWQQ<br>GQFWGQKPWSQHYHQGYY | 99,240 | 9.5 |

| | | | |
|---|---|---|---|
| U-CTD(ΔC16) | MHHHHHHHHKIEEGKLVIWINGDKGYNGLAEKFEKDTGIK<br>VTVEHPDKLEEKFPQVAATGDGPDIIFWAHDRFGGYAQSG<br>LLAEITPDKAFQDKLYPFTWDAVRYNGKLIAYPIAVEALS<br>LIYNKDLLPNPPKTWEEIPALDKELKAKGKSALMFNLQEP<br>YFTWPLIAADGGYAFKYENGKYDIKDVGVDNAGAKAGLTF<br>LVDLIKNKHMNADTDYSIAEAAFNKGETAMTINGPWAWSN<br>IDTSKVNYGVTVLPTFKGQPSKPFVGVLSAGINAASPNKE<br>LAKEFLENYLLTDEGLEAVNKDKPLGAVALKSYEEELAKD<br>PRIAATMENAQKGEIMPNIPQMSAFWYAVRTAVINAASGR<br>QTVDEALKDAQTNSSSVPGRGSSKKALPPEKKQNTGSKKS<br>NKNKSGKNQFNRGGGHRGRGGFNMRGGNFRGGAPGNRGGY<br>NRRGNMPQRGGGGGGSGGIGYPYPRAPVFPGRGSYSNRGN<br>YNRGGMPNRGNYNQNFRGRGNNRGYKNQSQGYNQWQQGQ | 83,770 | 9.5 |
| RGG1 | MHHHHHHHHKIEEGKLVIWINGDKGYNGLAEVGKKFEKDT<br>GIKVTVEHPDKLEEKFPQVAATGDGPDIIFWAHDRFGGYA<br>QSGLLAEITPDKAFQDKLYPFTWDAVRYNGKLIAYPIAVE<br>ALSLIYNKDLLPNPPKTWEEIPALDKELKAKGKSALMFNL<br>QEPYFTWPLIAADGGYAFKYENGKYDIKDVGVDNAGAKAG<br>LTFLVDLIKNKHMNADTDYSIAEAAFNKGETAMTINGPWA<br>WSNIDTSKVNYGVTVLPTFKGQPSKPFVGVLSAGINAASP<br>NKELAKEFLENYLLTDEGLEAVNKDKPLGAVALKSYEEEL<br>AKDPRIAATMENAQKGEIMPNIPQMSAFWYAVRTAVINAA<br>SGRQTVDEALKDAQTNSSSVPGRGSGGGRGGGHRGRGGFN<br>MRGGNFRGGAPGNRGGYNRRGNMPQRGGGGGG | 67,840 | 7.2 |
| RGG2 | MHHHHHHHHKIEEGKLVIWINGDKGYNGLAEVGKKFEKDT<br>GIKVTVEHPDKLEEKFPQVAATGDGPDIIFWAHDRFGGYA<br>QSGLLAEITPDKAFQDKLYPFTWDAVRYNGKLIAYPIAVE<br>ALSLIYNKDLLPNPPKTWEEIPALDKELKAKGKSALMFNL<br>QEPYFTWPLIAADGGYAFKYENGKYDIKDVGVDNAGAKAG<br>LTFLVDLIKNKHMNADTDYSIAEAAFNKGETAMTINGPWA<br>WSNIDTSKVNYGVTVLPTFKGQPSKPFVGVLSAGINAASP<br>NKELAKEFLENYLLTDEGLEAVNKDKPLGAVALKSYEEEL<br>AKDPRIAATMENAQKGEIMPNIPQMSAFWYAVRTAVINAA<br>SGRQTVDEALKDAQTNSSSVPGRGSGRGSYSNRGNYNRGG<br>MPNRGNYNQNFRGRGNNRGY | 72,310 | 6.6 |
| Δ-RGG1 | MHHHHHHHHKIEEGKLVIWINGDKGYNGLAEVGKKFEKDT<br>GIKVTVEHPDKLEEKFPQVAATGDGPDIIFWAHDRFGGYA<br>QSGLLAEITPDKAFQDKLYPFTWDAVRYNGKLIAYPIAVE<br>ALSLIYNKDLLPNPPKTWEEIPALDKELKAKGKSALMFNL<br>QEPYFTWPLIAADGGYAFKYENGKYDIKDVGVDNAGAKAG<br>LTFLVDLIKNKHMNADTDYSIAEAAFNKGETAMTINGPWA<br>WSNIDTSKVNYGVTVLPTFKGQPSKPFVGVLSAGINAASP<br>NKELAKEFLENYLLTDEGLEAVNKDKPLGAVALKSYEEEL<br>AKDPRIAATMENAQKGEIMPNIPQMSAFWYAVRTAVINAA<br>SGRQTVDEALKDAQTNSSSVPGRGSSKKALPPEKKQNTGS<br>KKSNKNKSGKNQFNSGGIGYPYPRAPVFPGRGSYSNRGNY<br>NRGGMPNRGNYNQNFRGRGNNRGYKNQSQGYNQWQQGQFW<br>GQKPWSQHYHQGYY | 97,750 | 9.1 |

| | | | |
|---|---|---|---|
| SGG1 | MHHHHHHHHKIEEGKLVIWINGDKGYNGLAEVGKKFEKDT<br>GIKVTVEHPDKLEEKFPQVAATGDGPDIIFWAHDRFGGYA<br>QSGLLAEITPDKAFQDKLYPFTWDAVRYNGKLIAYPIAVE<br>ALSLIYNKDLLPNPPKTWEEIPALDKELKAKGKSALMFNL<br>QEPYFTWPLIAADGGYAFKYENGKYDIKDVGVDNAGAKAG<br>LTFLVDLIKNKHMNADTDYSIAEAAFNKGETAMTINGPWA<br>WSNIDTSKVNYGVTVLPTFKGQPSKPFVGVLSAGINAASP<br>NKELAKEFLENYLLTDEGLEAVNKDKPLGAVALKSYEEEL<br>AKDPRIAATMENAQKGEIMPNIPQMSAFWYAVRTAVINAA<br>SGRQTVDEALKDAQTNSSSVPGRGSSKKALPPEKKQNTGS<br>KKSNKNKSGKNQFNSGGGHSGSGGFNMSGGNFSGGAPGNS<br>GGYNSSGNMPQSGGGGGGSGGIGYPYPRAPVFPGRGSYSN<br>RGNYNRGGMPNRGNYNQNFRGRGNNRGYKNQSQGYNQWQQ<br>GQFWGQKPWSQHYHQGYY | 99,240 | 9.1 |
| Δ-RGG2 | MHHHHHHHHKIEEGKLVIWINGDKGYNGLAEVGKKFEKDT<br>GIKVTVEHPDKLEEKFPQVAATGDGPDIIFWAHDRFGGYA<br>QSGLLAEITPDKAFQDKLYPFTWDAVRYNGKLIAYPIAVE<br>ALSLIYNKDLLPNPPKTWEEIPALDKELKAKGKSALMFNL<br>QEPYFTWPLIAADGGYAFKYENGKYDIKDVGVDNAGAKAG<br>LTFLVDLIKNKHMNADTDYSIAEAAFNKGETAMTINGPWA<br>WSNIDTSKVNYGVTVLPTFKGQPSKPFVGVLSAGINAASP<br>NKELAKEFLENYLLTDEGLEAVNKDKPLGAVALKSYEEEL<br>AKDPRIAATMENAQKGEIMPNIPQMSAFWYAVRTAVINAA<br>SGRQTVDEALKDAQTNSSSVPGRGSSKKALPPEKKQNTGS<br>KKSNKNKSGKNQFNRGGGHRGRGGFNMRGGNFRGGAPGNR<br>GGYNRRGNMPQRGGGGGGSGGIGYPYPRAPVFPKNQSQGY<br>NQWQQGQFWGQKPWSQHYHQGYY | 93,280 | 9.2 |
| SGG2 | MHHHHHHHHKIEEGKLVIWINGDKGYNGLAEVGKKFEKDT<br>GIKVTVEHPDKLEEKFPQVAATGDGPDIIFWAHDRFGGYA<br>QSGLLAEITPDKAFQDKLYPFTWDAVRYNGKLIAYPIAVE<br>ALSLIYNKDLLPNPPKTWEEIPALDKELKAKGKSALMFNL<br>QEPYFTWPLIAADGGYAFKYENGKYDIKDVGVDNAGAKAG<br>LTFLVDLIKNKHMNADTDYSIAEAAFNKGETAMTINGPWA<br>WSNIDTSKVNYGVTVLPTFKGQPSKPFVGVLSAGINAASP<br>NKELAKEFLENYLLTDEGLEAVNKDKPLGAVALKSYEEEL<br>AKDPRIAATMENAQKGEIMPNIPQMSAFWYAVRTAVINAA<br>SGRQTVDEALKDAQTNSSSVPGRGSSKKALPPEKKQNTGS<br>KKSNKNKSGKNQFNRGGGHRGRGGFNMRGGNFRGGAPGNR<br>GGYNRRGNMPQRGGGGGGSGGIGYPYPRAPVFPGSGSYSN<br>SGNYNSGGMPNSGNYNQNFSGSGNNSGYKNQSQGYNQWQQ<br>GQFWGQKPWSQHYHQGYY | 99,240 | 9.2 |
| SGG 1+2 | MHHHHHHHHKIEEGKLVIWINGDKGYNGLAEVGKKFEKDT<br>GIKVTVEHPDKLEEKFPQVAATGDGPDIIFWAHDRFGGYA<br>QSGLLAEITPDKAFQDKLYPFTWDAVRYNGKLIAYPIAVE<br>ALSLIYNKDLLPNPPKTWEEIPALDKELKAKGKSALMFNL<br>QEPYFTWPLIAADGGYAFKYENGKYDIKDVGVDNAGAKAG<br>LTFLVDLIKNKHMNADTDYSIAEAAFNKGETAMTINGPWA<br>WSNIDTSKVNYGVTVLPTFKGQPSKPFVGVLSAGINAASP<br>NKELAKEFLENYLLTDEGLEAVNKDKPLGAVALKSYEEEL<br>AKDPRIAATMENAQKGEIMPNIPQMSAFWYAVRTAVINAA<br>SGRQTVDEALKDAQTNSSSVPGRGSSKKALPPEKKQNTGS<br>KKSNKNKSGKNQFNSGGGHSGSGGFNMSGGNFSGGAPGNS<br>GGYNSSGNMPQSGGGGGGSGGIGYPYPSAPVFPGSGSYSN<br>SGNYNSGGMPNSGNYNQNFSGSGNNSGYKNQSQGYNQWQQ<br>GQFWGQKPWSQHYHQGYY | 99,240 | 7.8 |

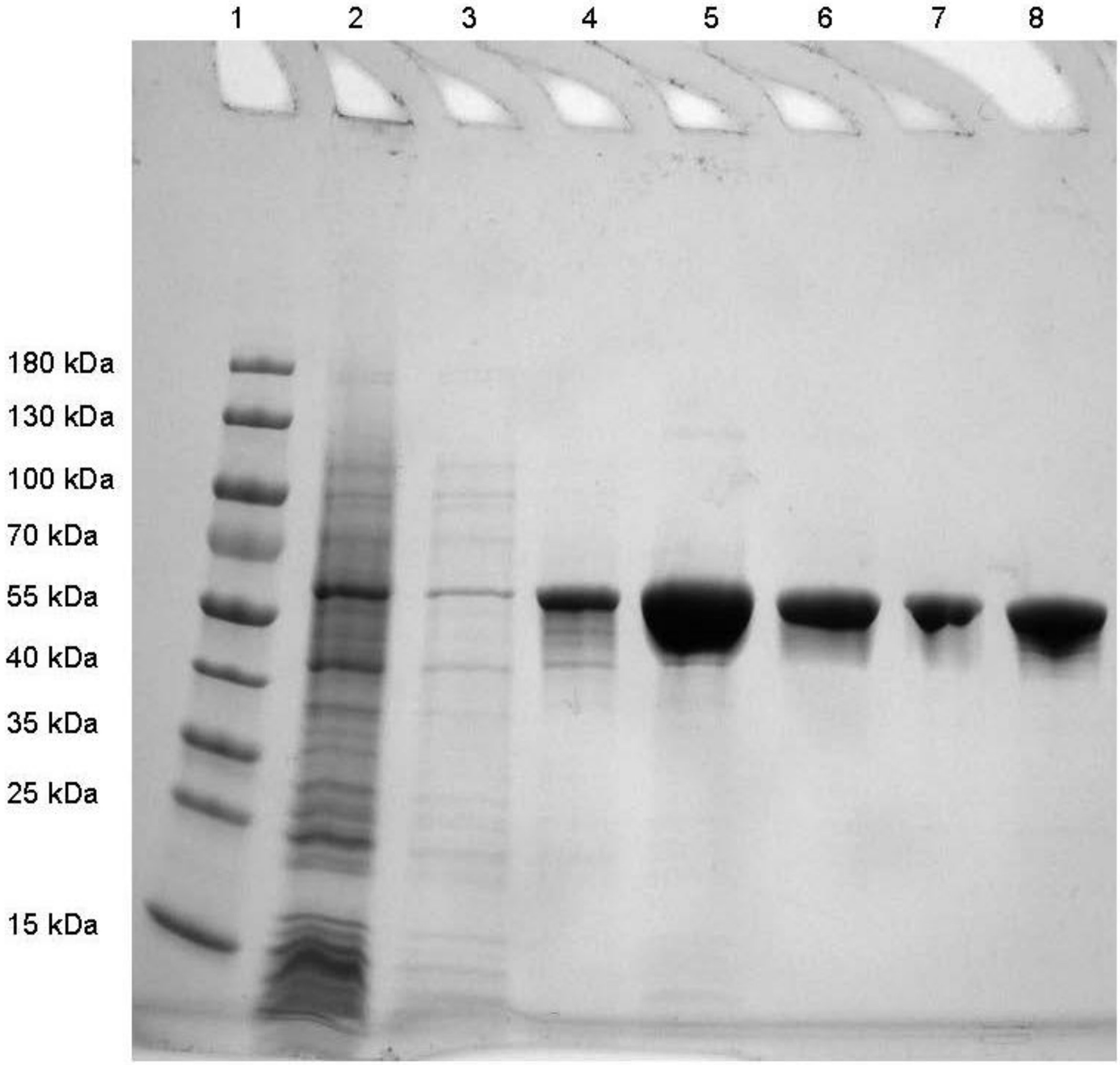

**Figure S1: SDS-PAGE analysis of protein purification.** Representative SDS-PAGE image of U-CTD purification. The protein was expressed and purified as described in the methods section. Lane 1: PAGERuler Pre-stained molecular weight marker. Lane 2: Flowthrough fraction of Ni purification. Lane 3: Wash 1 Fraction of Ni Purification. Lane 4: Wash 2 Fraction of Ni Purification. Lane 5: Elution fraction of Ni Purification. Lanes 6-8: SEC Fractions, which were pooled together to yield the final purified protein.

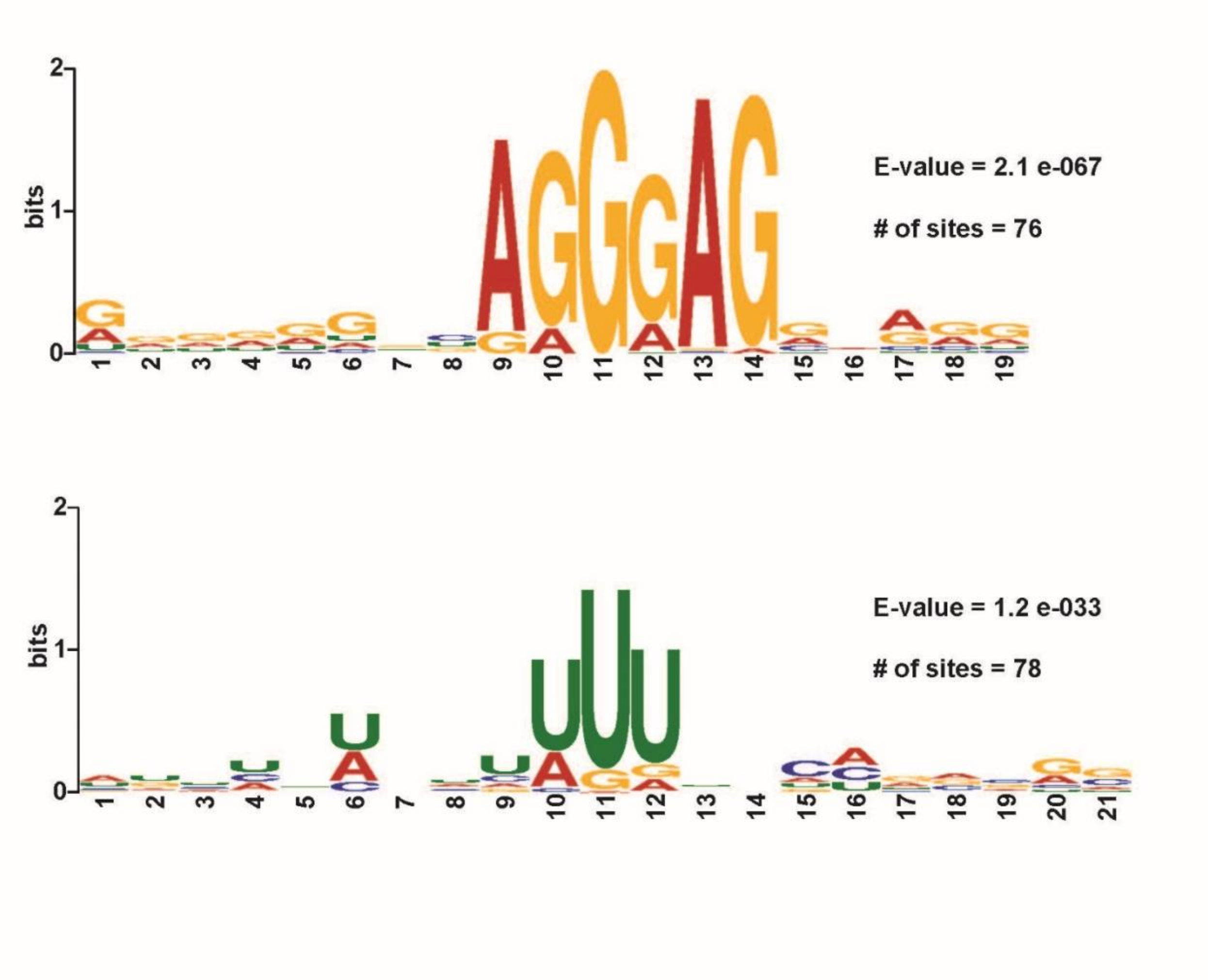

**Figure S2: MEME Motifs of the extended hnRNP U PureCLIP sites.** The final 323 hnRNP U PureCLIP sites were extended by 10 nucleotides in each and used as input for MEME motif detection, as in Figure 1. The top two motifs are shown.

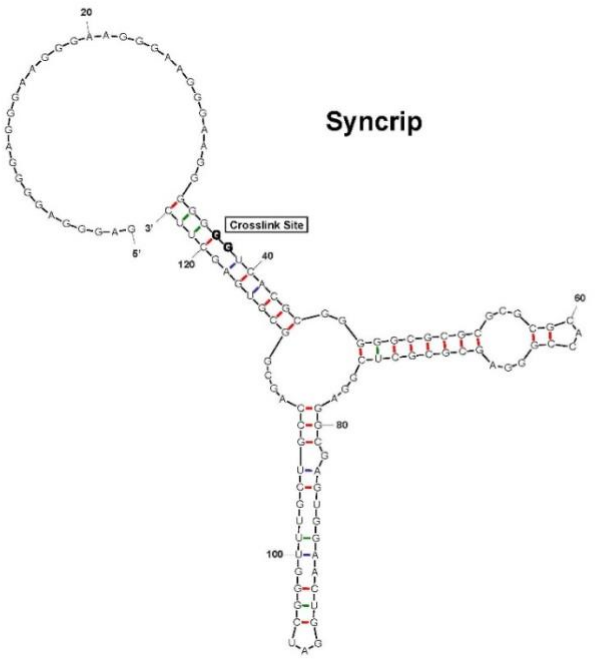
Syncrip
Crosslink Site

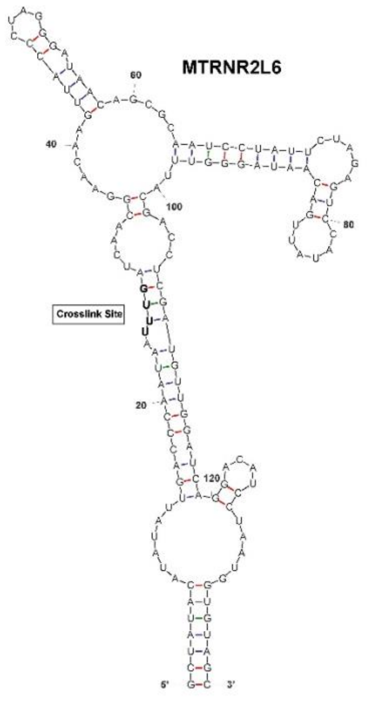
MTRNR2L6
Crosslink Site

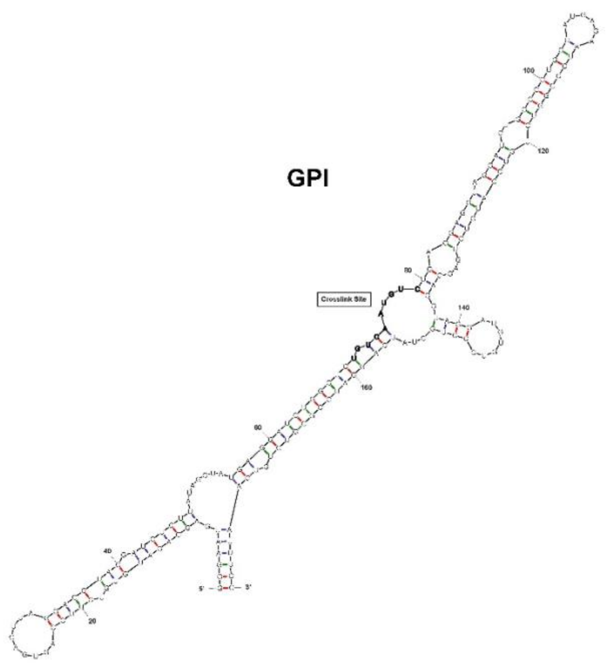
GPI
Crosslink Site

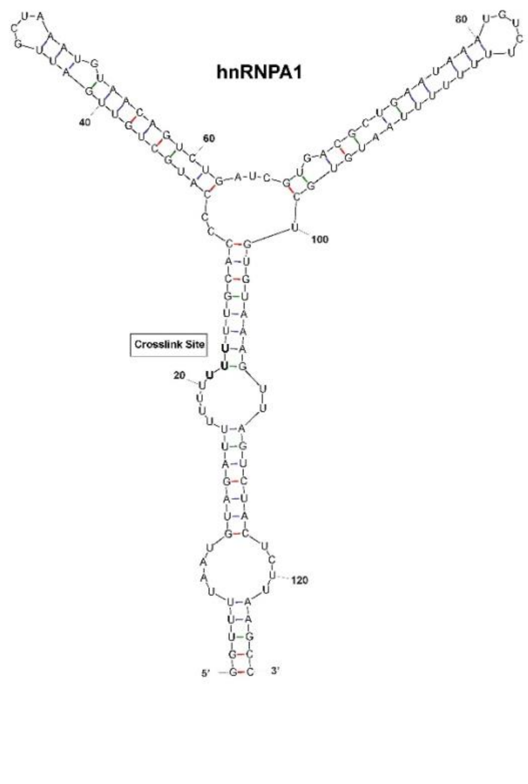
hnRNPA1
Crosslink Site

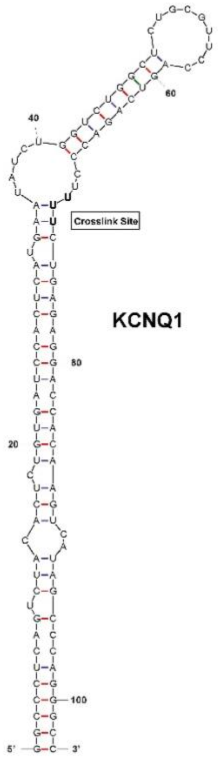
KCNQ1
Crosslink Site

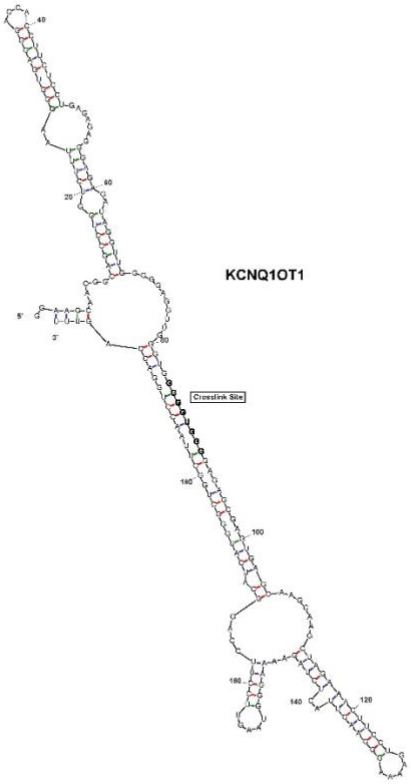
KCNQ1OT1
Crosslink Site

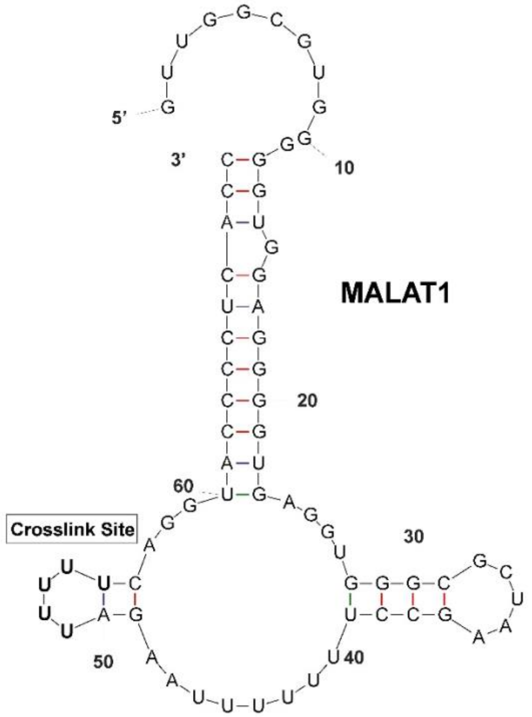
MALAT1
Crosslink Site

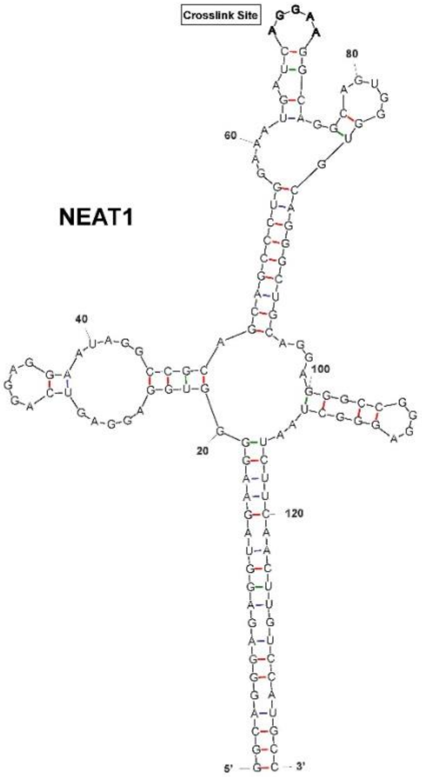
NEAT1
Crosslink Site

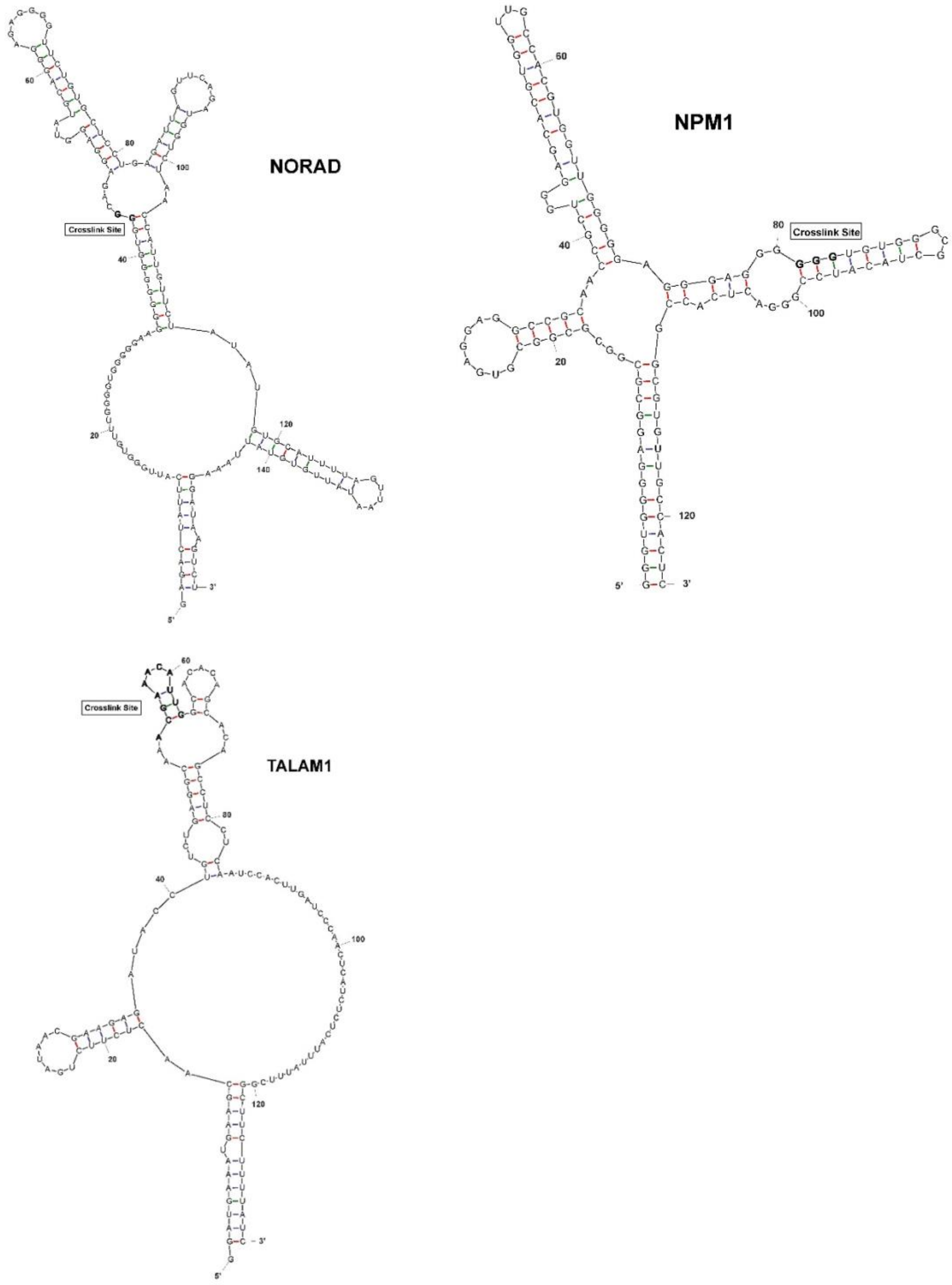

**Figure S3: Predicted secondary structures of survey RNAs.** RNA structures were predicted in Mfold using default settings, and the lowest MFE structure of each RNA was selected. The crosslink sites identified by PureCLIP are highlighted in bold lettering.

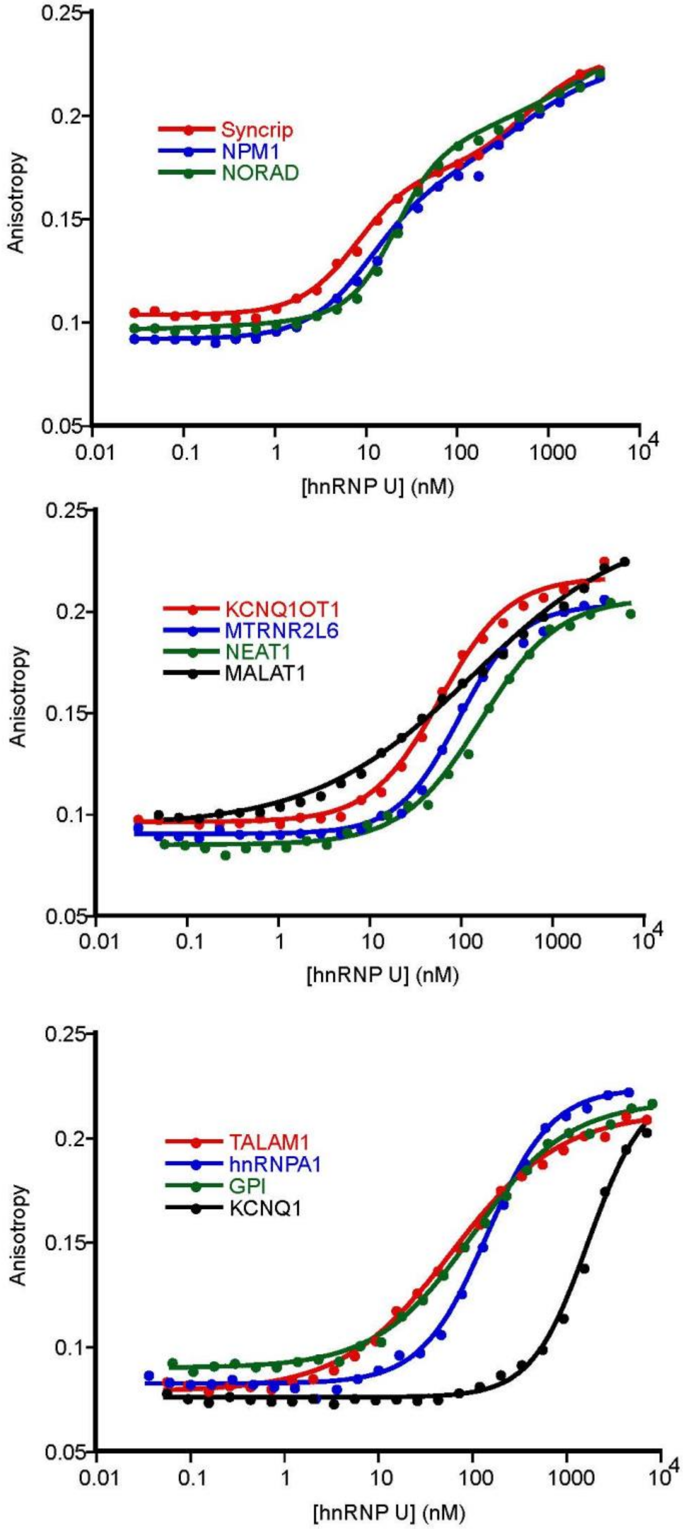

**Figure S4: Representative FA curves of hnRNP U bound to survey RNAs.** Each curve is a single replicate. NPM1, NORAD, and Syncrip curves were fit with a two-transition Hill equation. All other curves were fit with a one-transition Hill equation.

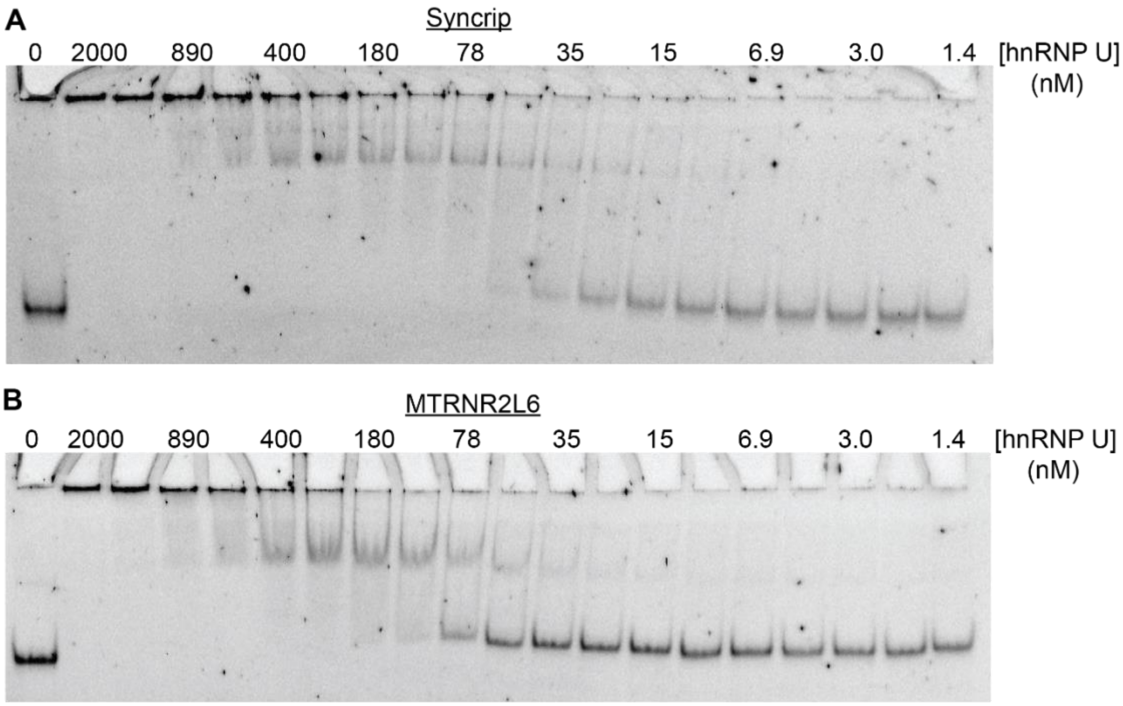

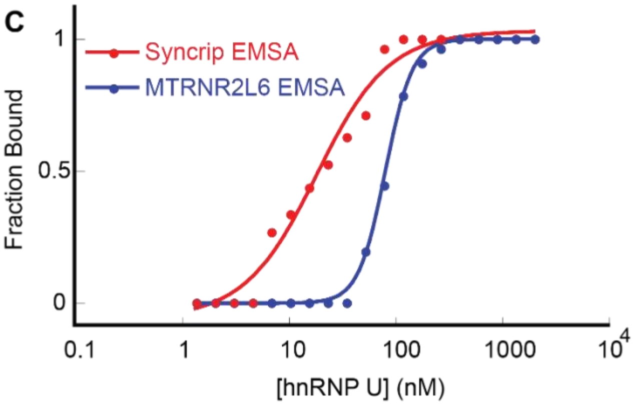

D

| RNA | $K_{Obs}$ (nM) | $N_H$ | $K_{Rel}$ (FA/EMSA) |
|---|---|---|---|
| Syncrip | 17 ± 4 | 1.2 ± 0.2 | 0.7 |
| MTRNR2L6 | 82 ± 1 | 3.4 ± 0.2 | 1.1 |

**Figure S5: Representative EMSAs of hnRNP U bound to survey RNAs.** (A) EMSA of hnRNP U bound to Syncrip RNA. (B) EMSA of hnRNP U bound to MTRNR2L6 RNA.(C) Normalized EMSA binding curves of hnRNP U bound to Syncrip and MTRNR2L6 RNAs. (D) Table of measured $K_{Obs}$, Hill coefficient, and relative $K_{Obs}$ between FA and EMSA measurements of hnRNP U bound to Syncrip and MTRNR2L6 RNAs.

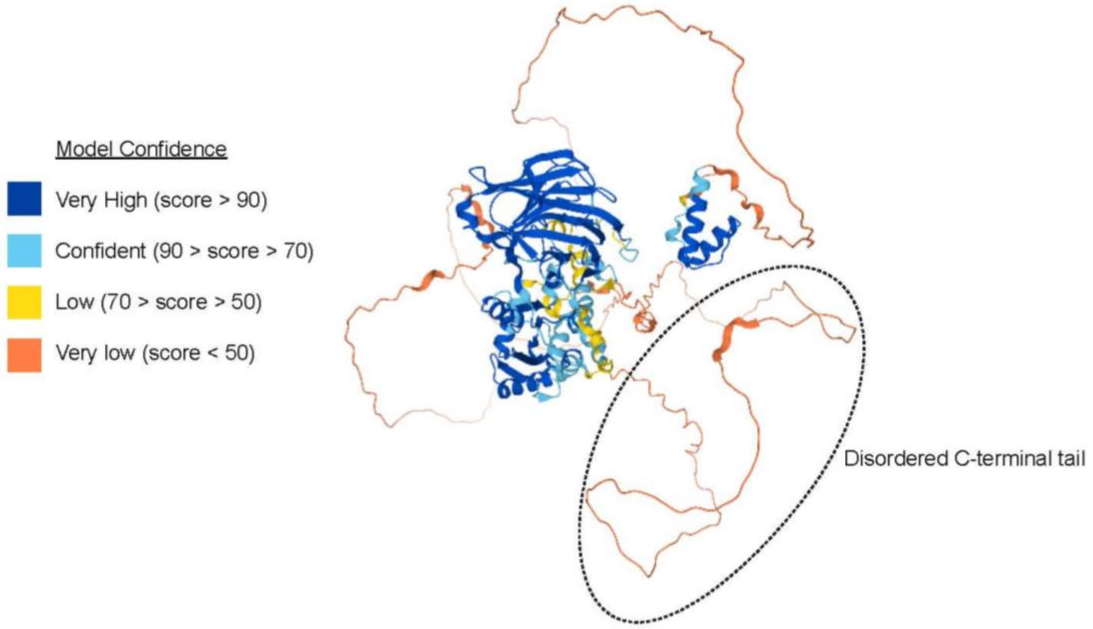

**Figure S6. Alphafold structure prediction of hnRNP U.** The structure of hnRNP U was predicted using Alphafold (1). The C-terminal tail containing the RNA-binding domain is highlighted in the dotted line.

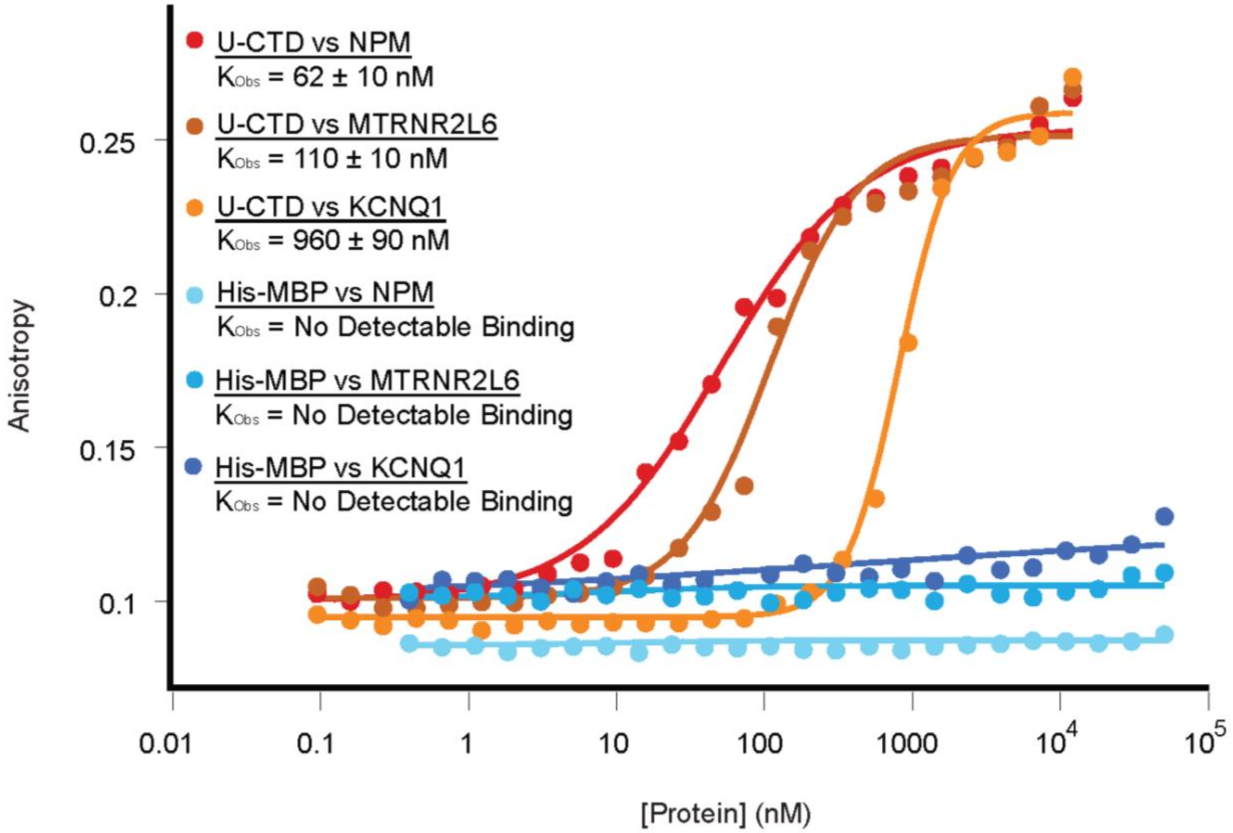

**Figure S7. MBP does not bind survey RNAs.** Representative FA curves of U-CTD and His-MBP bound to NPM1, MTRNR2L6, and KCNQ1 RNAs. These represent strong, intermediate, and weak targets of U-CTD.

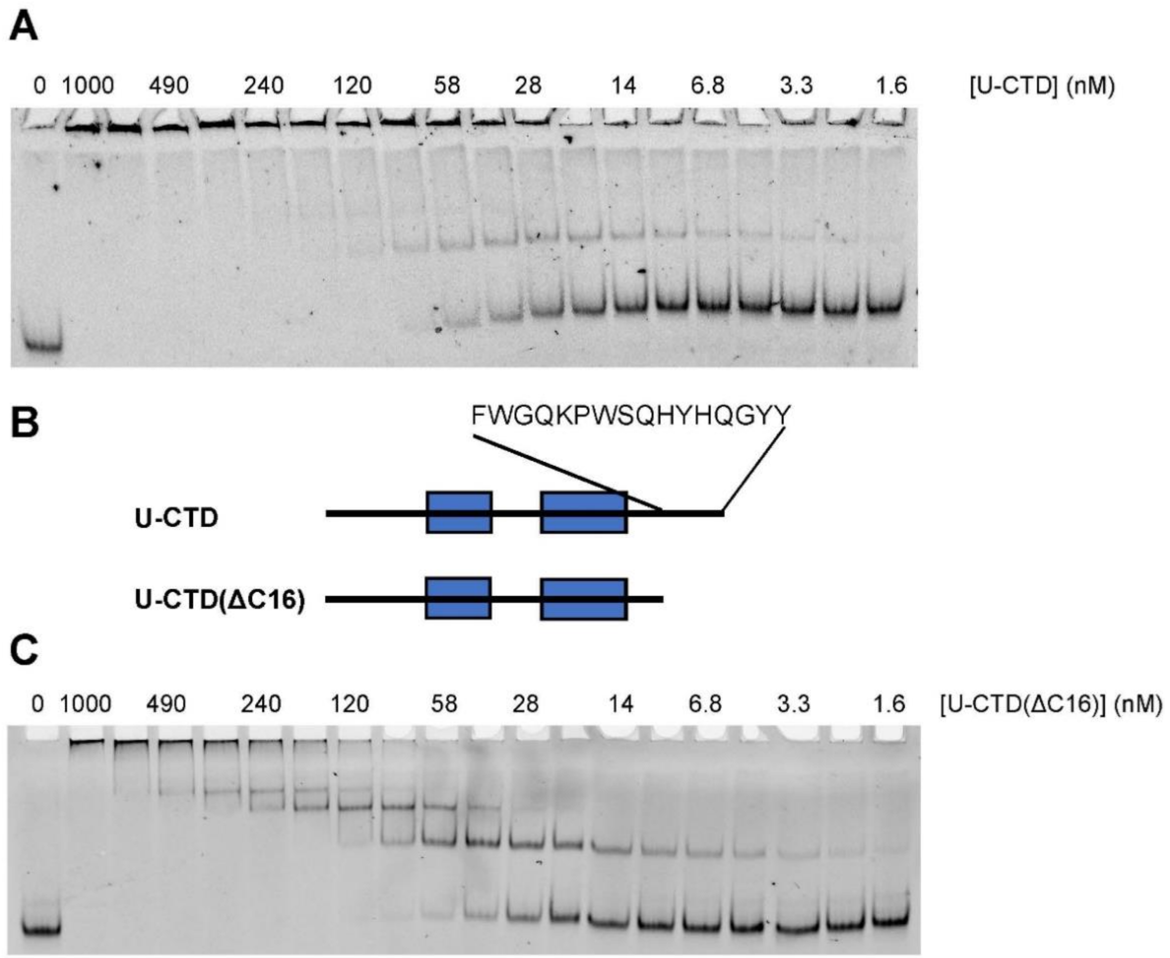

D

| RNA | $K_{Obs}$ (nM) U-CTD (ΔC16) | $N_{Hill}$ U-CTD (ΔC16) | $K_{Rel}$ U-CTD (ΔC16) / U-CTD |
|---|---|---|---|
| NPM1 | 95 ± 16 | 0.4 ± 0.1 | 1.5 |
| KCNQ1OT1 | 130 ± 10 | 1.0 ± 0.1 | 1.4 |
| NORAD | 140 ± 20 | 0.7 ± 0.1 | 1.5 |
| MTRNR2L6 | 140 ± 20 | 1.4 ± 0.1 | 1.3 |
| Syncrip | 150 ± 30 | 0.5 ± 0.1 | 1.8 |
| NEAT1 | 280 ± 10 | 1.3 ± 0.1 | 1.8 |
| GPI | 330 ± 10 | 1.6 ± 0.1 | 1.5 |
| TALAM1 | 330 ± 30 | 1.2 ± 0.1 | 2.3 |
| hnRNPA1 | 360 ± 10 | 1.4 ± 0.1 | 2.4 |
| MALAT1 | 880 ± 30 | 0.8 ± 0.1 | 3.1 |
| KCNQ1 | 1100 ± 100 | 2.3 ± 0.1 | 1.1 |

**Figure S8. Truncation of the C-terminus reduces well-shifting of protein-RNA complexes.** (A) EMSA of U-CTD bound to Syncrip RNA. (B) Schematic of U-CTD and the truncated U-CTD(ΔC16), with the amino acid sequence of the truncated region. (C) EMSA of U-CTD(ΔC16) bound to Syncrip RNA. (D) Table of average binding affinities between U-CTD(ΔC16) and the panel of survey RNAs as measured by FA, with associated SEM values. $K_{rel}$ shows a comparison with U-CTD. $N_{Hill}$ is the average Hill coefficient for each interaction.

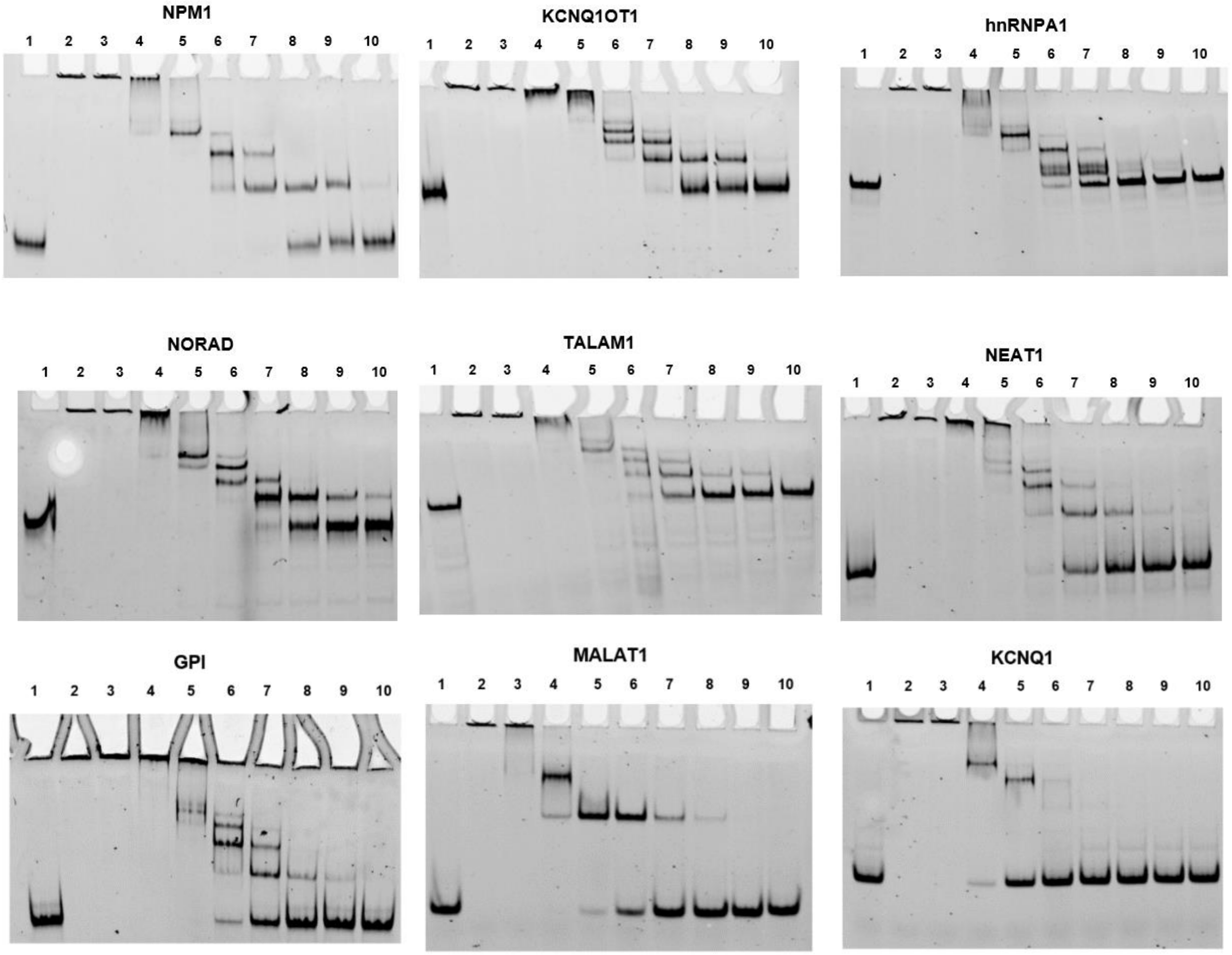

| Lane | [U-CTD(∆C16)] (nM) |
|---|---|
| 1 | 0 |
| 2 | 10000 |
| 3 | 3000 |
| 4 | 1000 |
| 5 | 300 |
| 6 | 100 |
| 7 | 30 |
| 8 | 10 |
| 9 | 3 |
| 10 | 1 |

**Figure S9: Electrophoretic Mobility Shift Assays (EMSAS) of survey RNAs.** Assays were performed as described in the methods section. U-CTD(∆C16) was used for all assays. Protein concentrations in each lane are indicated in the corresponding table.

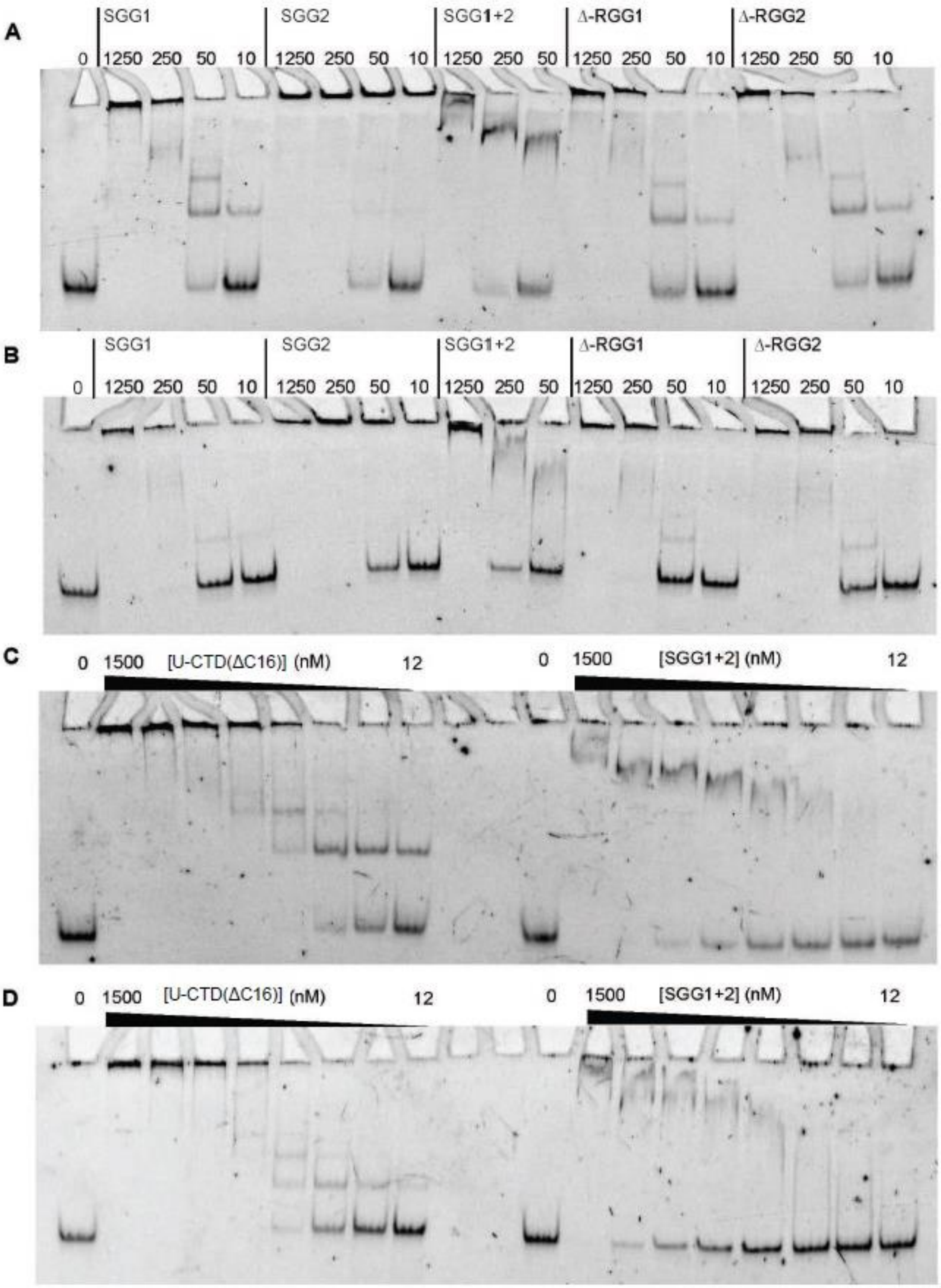

**Figure S10: EMSAs reveal altered binding modes by RGG/RG mutants.** (A) EMSA of RGG/RG domain variant constucts bound to Syncrip RNA. (B) EMSA of RGG/RG domain variant constucts bound to MTRNR2L6 RNA. (C) EMSA of U-CTD(ΔC16) and SGG1+2 bound to Syncrip RNA. (D) EMSA of U-CTD(ΔC16) and SGG1+2 bound to MTRNR2L6 RNA.
